# Demographic model and biological adaptation inferred from the genome-wide SNP data reveal tripartite origins of southernmost Chinese Huis

**DOI:** 10.1101/2021.10.18.464789

**Authors:** Guanglin He, Zhi-Quan Fan, Xing Zou, Xiaohui Deng, Hui-Yuan Yeh, Zheng Wang, Jing Liu, Quyi Xu, Ling Chen, Xiao-Hua Deng, Chuan-Chao Wang, Changhui Liu, Mengge Wang, Chao Liu

## Abstract

The culturally unique Sanya Hui (SYH) people are regarded as the descendants of ancient Cham people in Central Vietnam (CV) and exhibit a scenario of complex migration and admixture history, who were likely to first migrate from Central and South Asia (CSA) to CV and then to South Hainan and finally assimilated with indigenous populations and resided in the tropical island environments since then. A long-standing hypothesis posits that SYH derives from different genetic and cultural origins, which hypothesizes that SYH people are different from the genetically attested admixture history of northern Hui people possessing major Han-related ancestry and minor western Eurasian ancestry. However, the effect of the cultural admixture from CSA and East Asia (EA) on the genetic admixture of SYH people remains unclear. Here, we reported the first batch of genome-wide SNP data from 94 SYH people from Hainan and comprehensively characterized their genetic structure, origin, and admixture history. Our results found that SYH people were genetically different from the northern Chinese Hui people and harbored a close genomic affinity with indigenous Vietnamese but a distinct relationship with Cham, which confirmed the hypothesis of documented recent historical migration from CV and assimilation with Hainan indigenous people. The fitted admixture models and reconstructed demographic frameworks revealed an additional influx of CSA and EA ancestries during the historical period, consisting of the frequent cultural communication along the Southern Maritime Silk Road and extensive interaction with EA. Analyses focused on natural-selected signatures of SYH people revealed a similar pattern with mainland East Asians, which further confirmed the possibility of admixture-induced biological adaptation of island environments. Generally, three genetically attested ancestries from CV, EA, and CSA in modern SYH people supported their tripartite model of genomic origins.

## Introduction

The Hui people are a unique East Asian ethnolinguistic group predominantly comprising adherents of Islam and the second largest ethnic minority recognized by China with a population size of approximately 10.5 million (2010 census). The Hui people are mainly distributed in the northwestern provinces of China including Ningxia, Xinjiang, Gansu, and Qinghai, but communities also exist across the country (such as Henan and Hainan). The majority of Hui people speak Chinese while retaining some Arabic and Persian phrases (Dillon, 2013). And they also possess a unique cultural background, like other Chinese Muslim populations in Northwest China. The origin of Hui groups via demic diffusion (i.e., migration of people from non-East Asian regions), simple cultural diffusion (i.e., the indigenous East Asians converted to Hui people who were assimilated by Islam), or continuous migration (i.e., gene flow between people from non-East Asian regions and native East Asians) has been long debated (Chen et al., 2020; Liu et al., 2021a; Ma et al., 2021; Wang et al., 2019; Xie et al., 2019). Historical records have documented that Islam was first introduced into China by Arab merchants during the Tang dynasty about 1,400 years ago (ya). In several medieval dynasties (particularly the Tang, Song, and Yuan dynasties), traders, political emissaries, and soldiers continuously migrated from Arab, Persia, and Central and South Asia (CSA) to China mainly along with the land and maritime Silk Road (Gladney, 1998). The Huihui was the usual generic name for all Muslims in Imperial China (Gladney, 1996; Lipman, 1998). Nowadays, the term of Hui is applied by the Chinese government to indicate the only Sinophone Muslims, one of ten historically Islamic minorities (Lipman, 1998). Hence, the Hui people were believed to be of varied ancestries and likely to be the direct descendants of Silk Road immigrants who gradually mixed with indigenous East Asians (especially Han Chinese) (Chen, 1999; Gladney, 1998). However, it remains unclear the extent to which the Hui people had experienced a process of Chinesization and the non-East Asian ancestries had contributed to the gene pool of Hui people under the background of cultural admixture.

The genetic makeup of Hui people was investigated predominantly based on forensically relevant markers, such as autosomal short tandem repeats (A-STRs) (Deng et al., 2011; Yao et al., 2016), Insertion/Deletion polymorphisms (A-InDels) (Xie et al., 2018; Zhou et al., 2020; Zou et al., 2020), single nucleotide polymorphisms (A-SNPs) (He et al., 2018), X-chromosomal (Meng et al., 2014; Yang et al., 2017), Y-chromosomal (Guo, 2017; Wang et al., 2019; Xie et al., 2019; Zhao et al., 2017), and mitochondrial loci (Chen et al., 2020; Yao et al., 2004). These genetic analyses only revealed the forensic features of genetic markers in these culturally unique populations and demonstrated that northern East Asian Hui people descended from Central and West Asian migrants with Muslim cultural diffusion (Gladney, 1998). However, recent genetic findings demonstrated the close affinity between geographically different Hui groups and East Asian populations (Li et al., 2013; Yao et al., 2016; Zhou et al., 2020; Zou et al., 2020). He et al. sequenced 165 ancestry-informative SNPs (AISNPs) in Ningxia Huis and found the East Asian-dominant and European-minor ancestries in Hui people along the Silk Road (He et al., 2018), which highlighted the East Asian origin of the primary ancestors of Hui people. The matrilineal mtDNA profiles showed that Hui groups from northwestern China retain the minor genetic imprint of West Eurasian-specific lineages (Chen et al., 2020; Yao et al., 2004), while the patrilineal Y-DNA evidence identified relatively high levels of West Eurasian-related lineages in East Asian Hui groups (Wang et al., 2019; Xie et al., 2019), which indicated the sex-biased male-driven migrations. Furthermore, the genetic heterogeneity of Hui groups was confirmed via Y-SNP/Y-STR-based variations (Xie et al., 2019). It is remarkable, however, that the demographic history of the Hui groups has failed to be modeled because of the low resolution of those forensic-related markers and limited sample sizes. Thus, the admixture models and demographic processes of geographically different Hui people were needed to be further explored and characterized based on the denser genetic markers.

During the past decade, paleogenetic and genome-wide researches leveraging shared alleles and haplotypes transformed our knowledge of human population history (Elhaik et al., 2014; Wang, C.C. et al., 2021; Wang, T. et al., 2021; Wu et al., 2019), which provided many new insights into the demographic and evolutionary history of Chinese ethnolinguistic populations, including three ones focused on the formations of Hui people from central and northwestern China (Liu et al., 2021b; Ma et al., 2021; Wang et al., 2020a). Wang et al. explored the admixture history of Guizhou Huis and found that Guizhou Huis had ~6% West Eurasian-related ancestry and supported that cultural diffusion had played a pivotal role in the formation of the Hui people based on the array-based genome-wide SNPs (Wang et al., 2020b). A parallel study focused on Sichuan Huis also provided genetic evidence for the strong genetic affinity between Sichuan Huis and modern/ancient Northern East Asians and found that Sichuan Huis could be modeled as a mixture of major East Asian and minor West Eurasian (~6%) ancestries (Liu et al., 2021a). Recently, one pioneering research based on deep-whole-genome sequencing conducted by Xu’s group indicated that Ningxia Huis derived ~10% western ancestry and experienced an episode of two-date-two-wave admixture, the ancient admixture occurred ~1,025 ya and the recent admixture occurred ~500 ya (Ma et al., 2021). They also identified strong genetic affinity among Ningxia and Xinjiang Huis and Dungan people, consistent with their common pattern of migration and admixture processes (Ma et al., 2021). Additionally, these three genetic analyses have evidenced the existence of the sex-biased male-driven admixture in central and northwestern Chinese Hui people (Liu et al., 2021b; Ma et al., 2021; Wang et al., 2020a). In general, the genomic diversity of geographically different Hui groups implies the complex interplay between eastern and western Eurasians. Notably, most of these studies mainly focused on Hui groups from northern and southwestern China, the genetic structure of the Hui people from Hainan (the southernmost province of China) and their interaction with surrounding indigenous populations have been rarely explored.

Hainan Hui people also referred to as the Utsul people, are a Tsat-speaking (belonging to the Malayo-Polynesian group within the Austronesian language family) ethnic group which lives mainly in Sanya on the southernmost tip of Hainan Island. The Utsul group is officially recognized as the Hui nationality due to its Islamic faith, and they are actually ethnically, culturally, and linguistically distinct from other Hui Chinese. The Hainan Hui people are thought to be the descendants of Cham refugees who fled their homeland and moved to Hainan during the Ming Dynasty (Olson, 1998; Tran and Reid, 2006). However, some scholars argued that Cham refugees arrived on Hainan even earlier (in the Song dynasty) (Andaya, 2008; Grant and Sidwell, 2005). The close relatedness between the Hainan Huis and Cham was supported by their language affiliations (Edmondson and Gregerson, 1993; Thurgood, 2006). Whereas genetic evidence from uniparental markers revealed that Hainan Hui people showed a closer relationship with indigenous Hainan Li groups than with the Cham and other mainland Southeast Asian groups, which indicated that the formation of the Hainan Hui people likely involved massive assimilation of indigenous ethnic groups (Li et al., 2013). The estimated genetic makeup based on uniparental markers might bias our understanding of the admixture history of the Hainan Hui people. In this study, we generated high-density SNP data from 94 Hainan Hui people and comprehensively assessed the fine-scale admixture history of the Hui people by co-analyzing genome-wide data of publicly available modern and ancient Eurasian populations.

## Methods and materials

### Sample collection, genotyping and data merging

Salivary samples of 94 Hui individuals from Sanya in the south of Hainan were collected. Included participates were needed to be the direct offspring of indigenous Hui people with nonconsanguineous marriage with other populations in the last three generations. Our study was approved via the ethical recommendations of Xiamen University and followed the recommendations of the Helsinki Declaration. We obtained written informed consent from each of the participates. Infinium Global Screening Array (GSA) version 2 was used to genotype over 717K SNPs and quality-control was performed via PLINK (v1.90) (Chang et al., 2015) with the following parameters (mind: 0.01, geno: 0.01, --maf 0.01 and --hwe 10^-6^). Potentially existing relatives in three generations were tested via King software (Tinker and Mather, 1993). We merged our final dataset with previously published data genotyped via Illumina array (the merged Illumina dataset) (Chen et al., 2021; He et al., 2021; Liu et al., 2021b; Yao et al., 2021), and also combined it with the Affymetrix Human Origins dataset (the merged HO dataset) and the targeted-sequencing 1240K dataset (the merged 1240K dataset) from the Allen Ancient DNA Resource (AADR, https://reich.hms.harvard.edu/allen-ancient-dna-resource-aadr-downloadable-genotypes-present-day-and-ancient-dna-data). Other recently published modern and ancient genome-wide data from East Asia (EA) and Southeast Asia (SEA) were also collected (Mao et al., 2021; Wang, C.C. et al., 2021; Wang, T. et al., 2021; Yang et al., 2020).

### Frequency-based population genetic analysis

We used smartpca package in EIGENSOFT software (Patterson et al., 2006) to conduct principal component analysis (PCA) focused on the different scales of genetic variations of East Asians in different datasets. Ancient populations were projected and some potentially existing outliers were also included in the analysis (numoutlieriter: 0 and lsqproject: YES). Model-based clustering analyses were conducted using ADMIXTURE (V1.3.0) software (Alexander et al., 2009) based on the pruned dataset. We used PLINK (Chang et al., 2015) with these parameters (-indep-pairwise 200 25 0.4) to remove SNPs with strong linkage disequilibrium. We ran ADMIXTURE analysis 100 times and chose the best-fitted models based on the distribution of cross-validation error. Pairwise Fst genetic distances and corresponding distance matrixes were calculated using the in-house script based on the PLINK (Chang et al., 2015). We constructed the maximum-likelihood-based phylogenetic tree with different population split and migration events using TreeMix v.1.13 (Pickrell and Pritchard, 2012).

We further conducted a series of allele-sharing-based analyses using AdmixTools software (Patterson et al., 2012). We conducted pairwise qpWave analysis to test whether focused population pairs (left populations) formed one clade compared to the used outgroup sets (right populations) via the rank tests. We used eight worldwide populations (Mbuti, Ust_Ishim, Kostenki14, Papuan, Australian, Mixe, MA1, Jehai, and Tianyuan) as the right reference populations. We estimated the genetic affinity between SYH and other modern and ancient reference populations using qp3Pop (v435) package in the AdmixTools software (Patterson et al., 2012) via the tested form *f_3_*(reference, SYH; Mbuti). We also conducted admixture-*f_3_*-statistics in the form *f_3_*(source1 source2; SYH) to test the admixture signatures of SYH people, in which statistically negative *f_3_*-values with the Z-scores less than −3 denoted the admixture evidence with the two predefined ancestral surrogates. We conducted qpAdm-based admixture analysis using the same outgroup sets in the qpWave-based analysis via the AdmixTools package (Patterson et al., 2012). Two additional parameters were used in the qpAdm analysis (allsnps: YES; details: YES). To fit the phylogenetic framework with population divergence and admixture as well as estimate the genetic drift and admixture proportion of the involved population events, we used the R package of ADMIXTOOLS2 (Patterson et al., 2012) to explore the best fitted demographic models. We also used the ALDER 1.0 (Loh et al., 2013) to calculate the admixture times with two additional parameters: jackknife: YES and mindis: 0.005.

### Haplotype-based population genetic analysis

Dense SNP data can provide information on haplotype blocks, which are important for fine-scale population structure dissection. We used the Segmented HAPlotype Estimation & Imputation Tool (Shapeit v2) to phase the SNP data to their possible paternal and maternal haplotypes with the recommended parameters (--burn 10 --prune 10 --main 30) (Browning and Browning, 2011). Followingly, we used refined-ibd software (16May19.ad5.jar) and a block length of 0.1 to calculate pairwise IBD fragments (Browning and Browning, 2013). To paint the DNA block composition of the targeted chromosomes of SYH on the conditional of all possible sources from East Asians, Southeast Asians, and Central Asians, we ran ChromoPainterv2 software (Lawson et al., 2012) with all other sampled populations as the donor populations and SYH as the recipient population. We also used the R program of GLOBETROTTER (Hellenthal et al., 2014) with all sampled populations as their possible ancestral sources in the full analysis to identify, date, and describe admixture events of SYH people. We used fineSTRUCTURE (v4.0) (Lawson et al., 2012) to calculate the coancestry matrix and followingly conducted dendrogram clustering and PCA based on the haplotype data with default parameters (-s3iters 100000 -s4iters 50000 -s1minsnps 1000 -s1indfrac 0.1). Finally, we used R package of ReHH (Gautier et al., 2017) to calculate the cross-population extended haplotype homogeneity (XPEHH) with southern Han Chinese and northern Han Chinese as the reference populations. We also used the ReHH to calculate the integrated haplotype score (iHS) and used the online tool of Metascape (Zhou et al., 2019) to perform enrichment analysis.

## Results

### The patterns of population diversity and genetic structure

We genotyped over 700K genome-wide SNPs in 94 self-identified Hui people from Sanya in Hainan province, Southmost of mainland China. We estimated the kinship and removed six individuals within three generations and remained the final genome-wide SNP data of 88 Hui people in the following population genetic analysis. To comprehensively explore the population structure of Hui people and deeply characterize the genetic relationship between Hui people and reference populations from Eurasia, we merged our newly-genotyped data with three different datasets with different SNP densities: the merged Illumina (533943 SNPs, 533K), HO (56946 SNPs, 56K) and 1240K (158661 SNPs, 158K) datasets. We firstly applied PCA analysis to obtain the overall genetic structure and the potential relationship among the included modern and ancient eastern Eurasian samples in the merged HO dataset (**Figure 1A**), which possessed more geographically different populations but with low-density SNPs. PCA based on genetic diversity of eastern Eurasians highlighted two genetic clines respectively from northern and southern EA. We found SYH people were localized between these two clines and separated from these two clines (**Figure 1B**). To further explore the genetic relationship between SYH and southern East Asians and Southeast Asians, we focused on the genetic diversity of Sinitic people and populations belonged to the five language families in southern eastern Eurasia (Austronesian, Austroasiatic, Tai-Kadai, Hmong-Mien, and Tibeto-Burman). PCA results showed three directions: Hong-Mien and Austronesian (Kankanaey and Ami) people localized in the top, Sinitic and Tibeto-Burman (Bai_Heqing, Naxi_HGDP, and Pumi_Lanping) people localized in the lower left, and other Austronesian, Austroasiatic, Tibeto-Burman and Tai-Kadai people localized in the lower right. Studied Hui people also localized among the three clines or clusters and had a close relationship with some Austroasiatic people (Kinh and Vietnamese) (**Figure 1C**). To confirm the genetic similarities and differences between Hui and Han Chinese populations, we further explored the genetic relationship between Huis and Hans, as well as among geographically different Hui people (Sanya, Guizhou, and Sichuan). PCA restricted to the genetic variations of studied and Sinitic-speaking populations showed that SYH had a unique genetic structure not only distinguishable from Han Chinese but also from other geographically diverse Hui people (**Figure 1D**). To further directly present the unique genetic structure of SYH people, we conducted model-based ADMIXTURE analysis among the modern populations included in the merged HO dataset and chose the best-fitted models based on the lowest cross-validation error value (K=9). Ancestry composition inferred from ADMIXTURE showed SYH possessed a specific ancestry maximized in themselves, which also existed in Austroasiatic, Austronesian, and Tai-Kadai people in Vietnam (**Figure 1E**).

**Figure 1.**
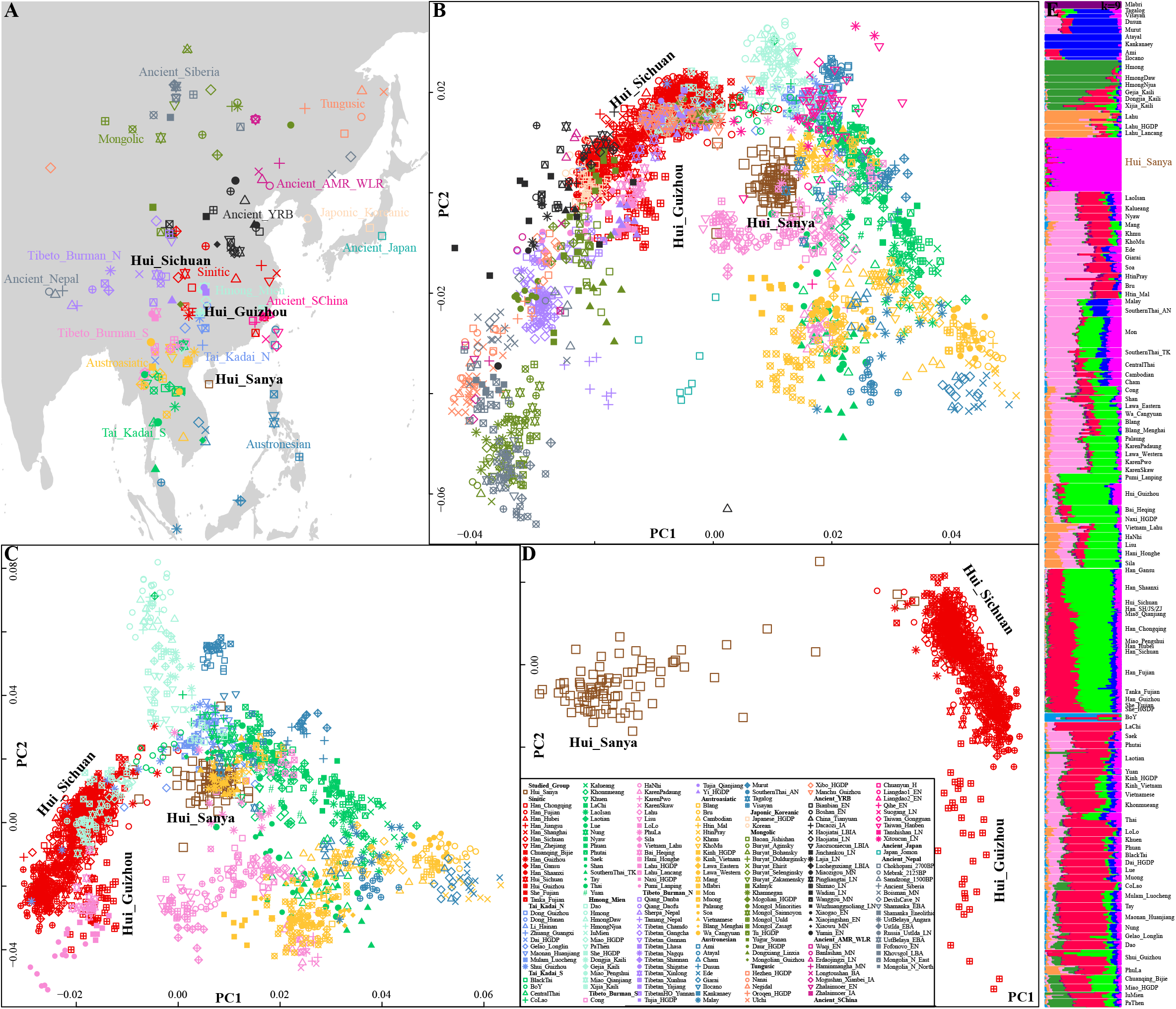
General population structure of Sanya Hui people and modern/ancient East Asians based on the merged HO dataset. (**A**), the geographical position of Sanya Hui people and other 229 included reference populations. Modern populations were color-coded via the language categories and ancient populations were colored via geographical positions. (**B**), Principal component analysis (PCA) among 230 modern and ancient East Asian populations, in which modern Mlabri and ancient populations were projected. (**C**), PCA results among 104 populations from South China and Southeast Asia. (**D**), Patterns of genetic relationship between Hui and Han Chinese populations, which included three Hui populations and 13 Han Chinese populations. (**E**), Model-based ADMIXTURE results showed the ancestry composition with the lowest cross-validation error (K=9).

To explore whether there is a deviation toward western Eurasians of SYH in the PCA and further validate the identified genetic similarity in the HO dataset, we then performed PCAs based on the merged Illumina and 1240K datasets. PCA based on East Asians and Central/South Asians showed that East Asians, including SYH, were separated from other western Eurasian sources (**Figure 2A**) along PC1. We could observe that Mongolic-speaking Mongolians in Xinjiang, Tibeto-Burman-speaking Tus and Guizhou Huis deviated toward Central Asian populations, consistent with reported western Eurasian affinity (Zhao et al., 2020). However, SYH people were located at the end of the East Asian north-to-south genetic clines and separated from other Chinese populations. We further confirmed their isolated genetic position among modern and ancient East Asians based on the merged 1240K (**Figures 2B~C**). To directly validate the consistency of population genetic relationship, we also explore the genetic relationship between Hui and Han Chinese based on the merged Illumina dataset, and we found that the similar genetic patterns of genetic relationships among geographically different Huis and Hans could be visualized via both lower-density and higher-density datasets (**Figures 2D**), which suggested that the population genetic findings based on the merged HO dataset are robust. Subsequently, we also performed PCA analysis focused on the three Hui populations (Sanya, Guizhou, and Sichuan) and we identified obvious population differentiation, suggesting their different evolutionary and admixture history (**Figure 2E**). As expected, ADMIXTURE results based on modern and ancient populations in the merged 1240K dataset also identified one Hui-specific ancestry (orange ancestry), which was dominant in SYH people and also existed in Austroasiatic people of Blang and Wa (**Figure 2F**). We also found an ancestry related to ancient Fujian and Taiwan people who had contributed some genetic materials to SYH people, however, some SYH individuals shared more ancestry with Hans and showed as outliers in the PCA analysis, which might be the recently admixed individuals deriving ancestral component from Hans and removed in the following population genetic analysis focusing on the genetic origin, admixture history and biological adaptation of SYH people.

**Figure 2.**
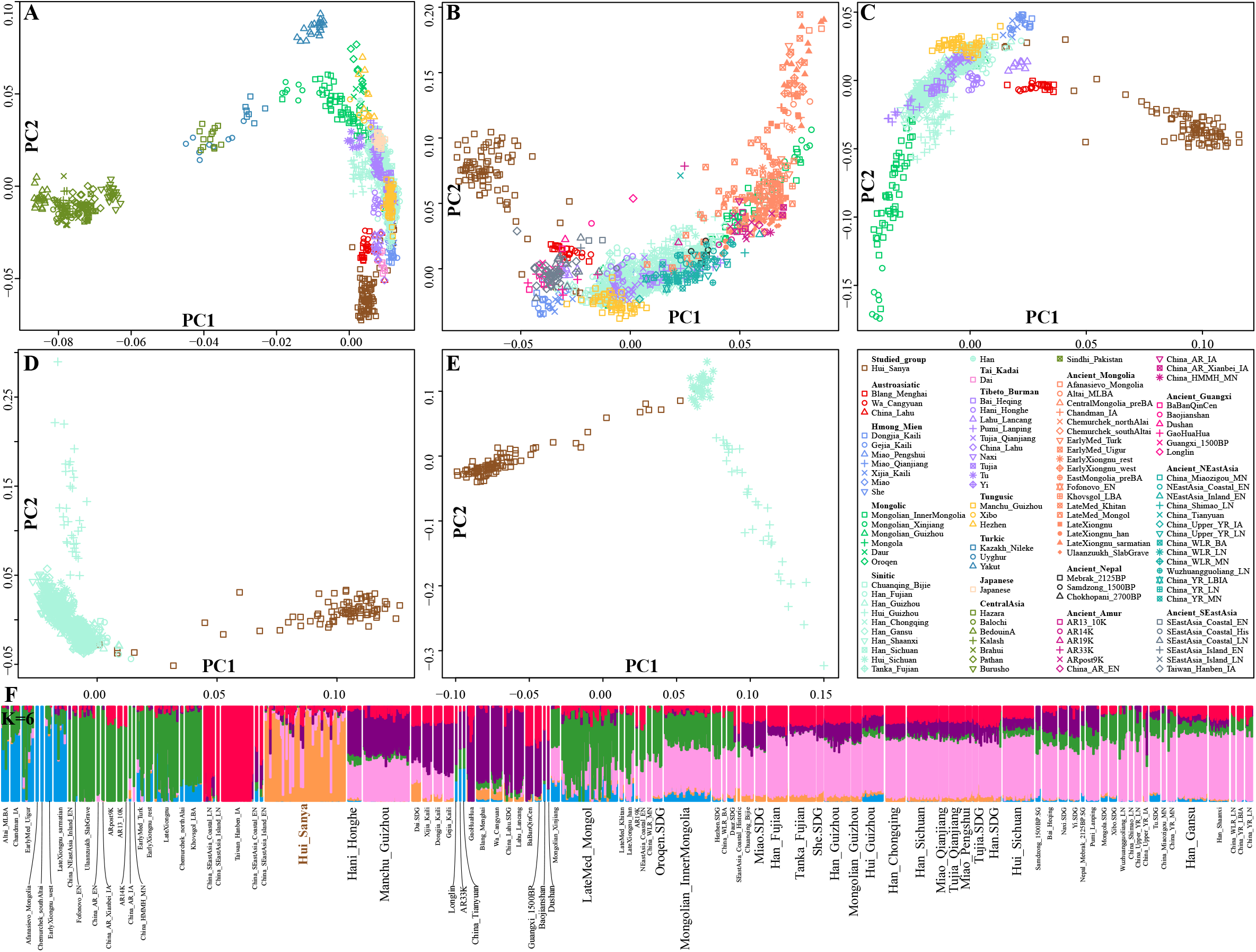
Genetic structure of central and eastern Eurasian populations based on denser SNP data. (**A**), PCA patterns among 54 modern populations from Central and East Asians. (**B~C**), PCA results of East Asian populations with (**B**) and without (**C**) ancient populations from the Mongolian Plateau based on the merged 1240K dataset. (**D**), Genetic relationship between Hui and Chinese populations based on the Illumina dataset. (**E**) Genetic relationship among three geographically different Hui populations. (**F**), ADMIXTURE results with six predefined ancestral sources among populations in the merged 1240K dataset.

### The genomic affinity between Sanya Huis and Eurasians

We first assessed the genetic affinity between Hui people and modern and ancient Eurasian reference populations via pairwise Fst genetic distances. The estimated affinity among 24 Chinese populations based on the Illumina dataset showed that SYH possessed the lowest Fst values with Guizhou Han (0.0110), followed by Miaos and Hans (**Table S1**). It is interesting to observe that SYH had a distant relationship with Sichuan Hui (0.0134) and Guizhou Hui (0.0184), consistent with the clustering patterns in the PCA and ADMIXTURE. When more populations from non-EA were included based on the merged HO dataset (**Table S2**), we identified the strongest genetic affinity between SYH and Austroasiatic people from Vietnam (Kinh: 0.0094 and Vietnamese: 0.0097), followed by Tai-Kadai people in South China, which provided clues for genetic contribution from Vietnam and South China into SYH people. Based on the shared genetic drift estimated via outgroup-*f_3_*(Reference populations, SYH; Mbuti) (**Table S3**), we further identified a close genetic affinity with Austronesian (Kankanaey: 0.3015) and Hmong-Mien speakers (0.3011 for Dao, Ami, and Gejia), suggesting the massive intrafraction among them. We also assessed the genetic relationship between SYH and ancient East Asians and we found SYH had the closest genetic relationship with the Iron Age population (Luoheguxiang_LBIA: 0.2969) and historical Guangxi population (GaoHuaHua: 0.2967).

Formal tests based on the admixture-*f_3_*-statistics showed that no statistically significant negative *f_3_*-values were observed in *f_3_*(source1, source2; SYH) among modern and ancient ancestral surrogates (**Table S4**). Thus, to further explore the possible asymmetrical gene flow events between SYH and other modern and ancient reference populations, we conducted a series of *f_4_*-statistics (**Tables S5~7**). First, we conducted *f_4_*(Hui1, Hui2; reference, Mbuti) to explore the genetic relationship between SYH and northern Chinese Hui people from Guizhou and Sichuan (**Table S5**). The lowest *f_4_*-values were observed in the *f_4_*(SYH, Hui_Sichuan; China_Upper_YR_LN, Mbuti)=-19.365*SE, followed by other ancient reference populations from the Yellow River Basin, Mongolian Plateau, southern Siberia and Tibetan Plateau and modern northern East Asians. Similar patterns were also identified focused on Guizhou Hui in the *f_4_*(SYH, Hui_Guizhou; reference, Mbuti), which suggested that Sichuan and Guizhou Hui people harbored more northern East Asian ancestry compared with SYH. However, statistically significant positive *f_4_*-values were observed in *f_4_*(SYH, Hui_Guizhou; Fujian/Guangxi ancients and Hmong-Mien people, Mbuti), which suggested that SYH people harbored more southern East Asian ancestry compared with Guizhou Hui. We only observed one statistically positive Z-score focused on Sichuan Hui in *f_4_*(SYH, Hui_Sichuan; Guangxi_1500BP, Mbuti)=3.699*SE. We conducted symmetrically *f_4_*-statistics *f_4_*(reference1, referene2; SYH, Mbuti) and confirmed that SYH people also shared more alleles with Han Chinese populations relative to most of the non-Han Chinese reference populations.

We then focused on the genetic relationship between Cham and SYH people using *f_4_*-statistics (**Tables S6~7**). SYH people might be the descendants of ancient Cham people based on the historic documents. If SYH people were the direct decedents of Cham people with no additional gene flow events, we expected to observe statistically negative *f_4_*-values in *f_4_*(reference, SYH; Cham, Mbuti) and non-statistically significant values in *f_4_*(Cham, SYH; reference, Mbuti). However, we only observed statistically negative *f_4_*-values in *f_4_*(Reference populations with western Eurasian ancestry, SYH; Cham, Mbuti) and statistically positive *f_4_*-values in *f_4_*(Tai-Kadai, SYH; Cham, Mbuti), which suggested that Cham shared more alleles with SYH people only relative to western Eurasian-related populations and with modern Tai-Kadai people compared with SYH people. Further results from *f_4_*(Cham, SYH; reference, Mbuti) did not show non-significant *f_4_*-values but harbored more significant negative *f_4_*-values, which suggested that other used reference populations from EA and SEA contributed more ancestry into Hui’s gene pool compared with Cham. It also suggested that Cham possessed a relatively distant genetic relationship with East Asians.

As a close genetic affinity was identified between Kinh and SYH people, we also assumed Kinh as the direct ancestor of SYH people and tested this hypothesis using the aforementioned *f_4_*-statistics (**Table S7**). More statistically negative *f_4_*-values in *f_4_*(Western Eurasian/Austronesian, Austroasiatic and Tibeto-Burman in SEA, SYH; Kinh, Mbuti) showed a genetic affinity between Kinh and Hui people. The lowest *f_4_*-values in *f_4_*(Kinh_HGDP, SYH; Bedouin, Mbuti)=-2.081*SE showed marginally additional gene influx from Arab populations related to Bedouin into Hui people compared with Kinh. The estimated positive in *f_4_*-statistics also showed that we can identified the obvious northern East Asian gene flow into Kinh via *f_4_*(Kinh_HGDP, Hui; East Asians, Mbuti).

MtDNA evidence has demonstrated that SYH people were assimilated with indigenous people and had the closest genetic affinity with geographically close Lis (Li et al., 2013). Thus, we further explored their genetic relationship via *f_4_*-statistics (**Table S7**). Results from *f_4_*(southern and central Tai/Karen/LaoIsan/Sila/HaNhi/Shan/Lisu/Hani, SYH; Li_Hainan, Mbuti) showed statistically negative values, which suggested a close genetic relationship between SYH and Li people compared with Southeast Asians. However, the estimated positive values in *f_4_*(Li_Hainan, SYH; Hmong-Mien/Tai-Kadai, Mbuti) showed that Hainan Li shared more alleles with Tai-Kadai and Hmong-Mien people compared with SYH, which suggested the differentiated genetic history between Hui and Li people in Hainan province. This pattern may be caused by the influence of the identified western Eurasian ancestry in SYH people.

Additionally, we conducted pairwise qpWave analysis (**Figure 3**) among 124 modern populations from China and SEA and five ancient populations from Guangxi province (**Table S8**). Heatmap among 94 populations (except Tibeto-Burman) showed that SYH people clustered closely with some of Austronesian, Austroasiatic, and Tai-Kadai people from Vietnam and showed a relatively heterogeneous relationship with northern Han Chinese populations, suggesting that major additional genetic admixture with southern Chinese indigenes rather than with Han people contributed more to the formation of SYH. Focused on the individual pairwise tests, we found evidence for genetic homogeneity between SYH people with 1500-year-old Guangxi BaBanQinCen, five Austroasiatic (Palaung, Kinh, Muong, Western Lawa and Blang), two Austronesian (Atayal and Ilocano), one Hmong-Mien (Njua Hmong), two Tai-Kadai (Khonmueang and Shan) and seven southern Tibeto-Burman (Lahu, Cong, Sila, PhuLa, HaNhi, Lisu and KarenPwo), which suggested that the East Asian ancestor of SYH people was related to indigenous people from South China and SEA during the extensive population migration and admixture.

**Figure 3.**
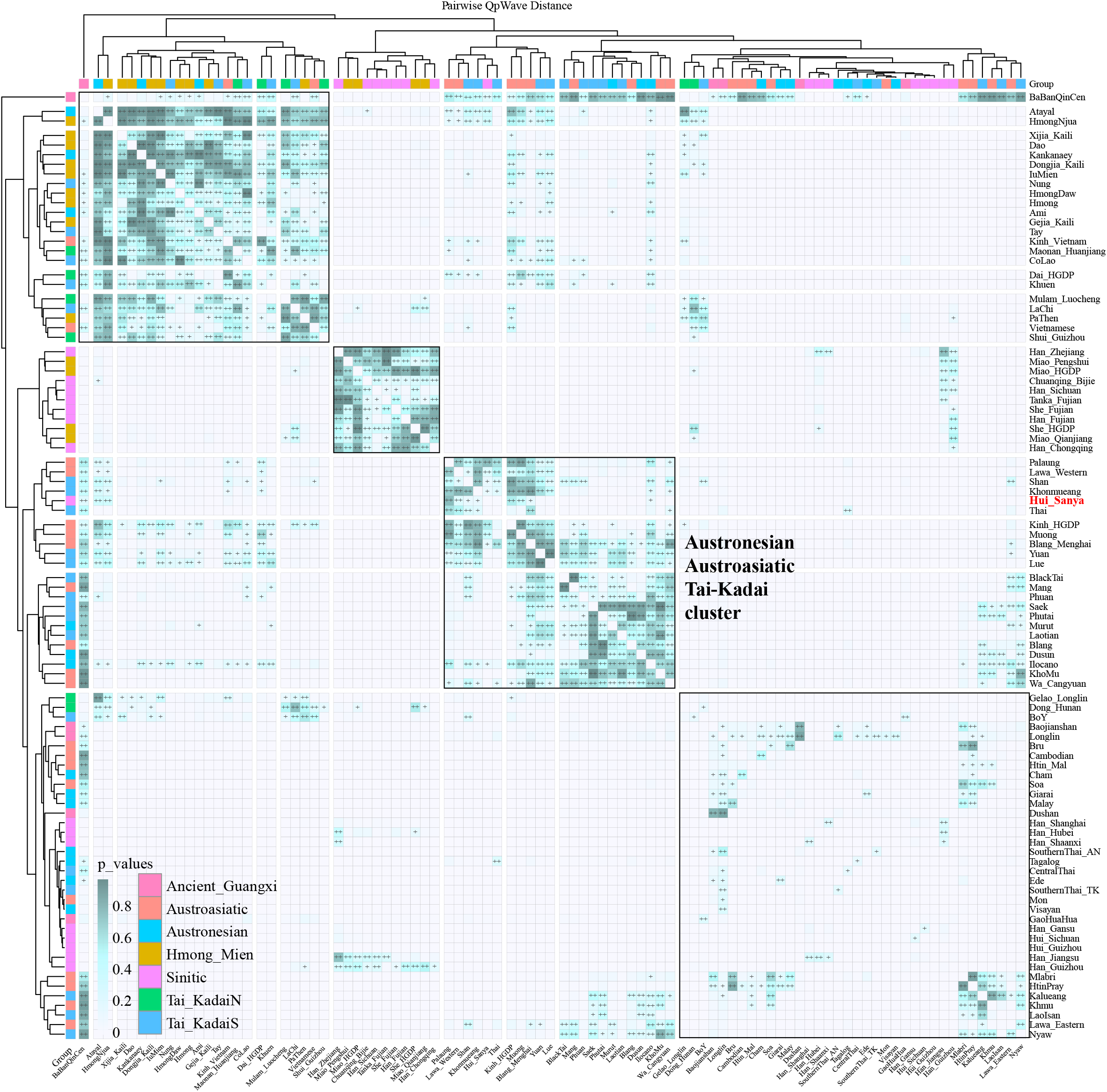
Pairwise qpWave analysis among 89 modern and 5 Guangxi ancient populations. p values larger than 0.05 were marked as “++” and p values ranging from 0.01 to 0.05 were marked as “+”. Populations with genetic homogeneity were trended to be clustered together.

### Admixture modeling of SYH people

To explore the basic phylogenetic position of SYH with Eurasian reference populations, we first constructed the TreeMix-based phylogenetic tree with population splits and gene flow events among 24 Chinese populations (**Figure 4A**). We identified three major branches respectively associated with Hmong-Mien, Austroasiatic and Sinitic people and SYH clustered with the Austroasiatic branch. However, Hui people from Guizhou and Sichuan clustered with Sino-Tibetan people. We observed one gene flow from Hani_Honghe into Hui_Guizhou and no obvious gene flow from other people into SYH people was observed. Based on the merged 1240K dataset among 53 populations (**Figure 4B**), we found that SYH people clustered with Chinese Tibeto-Burman, Austroasiatic, and Tai-Kadai people from Yunnan and Hmong-Mien people from Guizhou, which simultaneously obtained the gene flow from western Eurasian Balochi with a proportion of 0.07±0.06 (p_value: 0.0025). In the third TreeMix-based phylogenetic tree (**Figure 4C**), SYH first clustered closely with Kinh and Vietnamese from Vietnam and then with Tai-Kadai and Hmong-Mien people, which also absorbed one western Eurasian gene flow.

**Figure 4.**
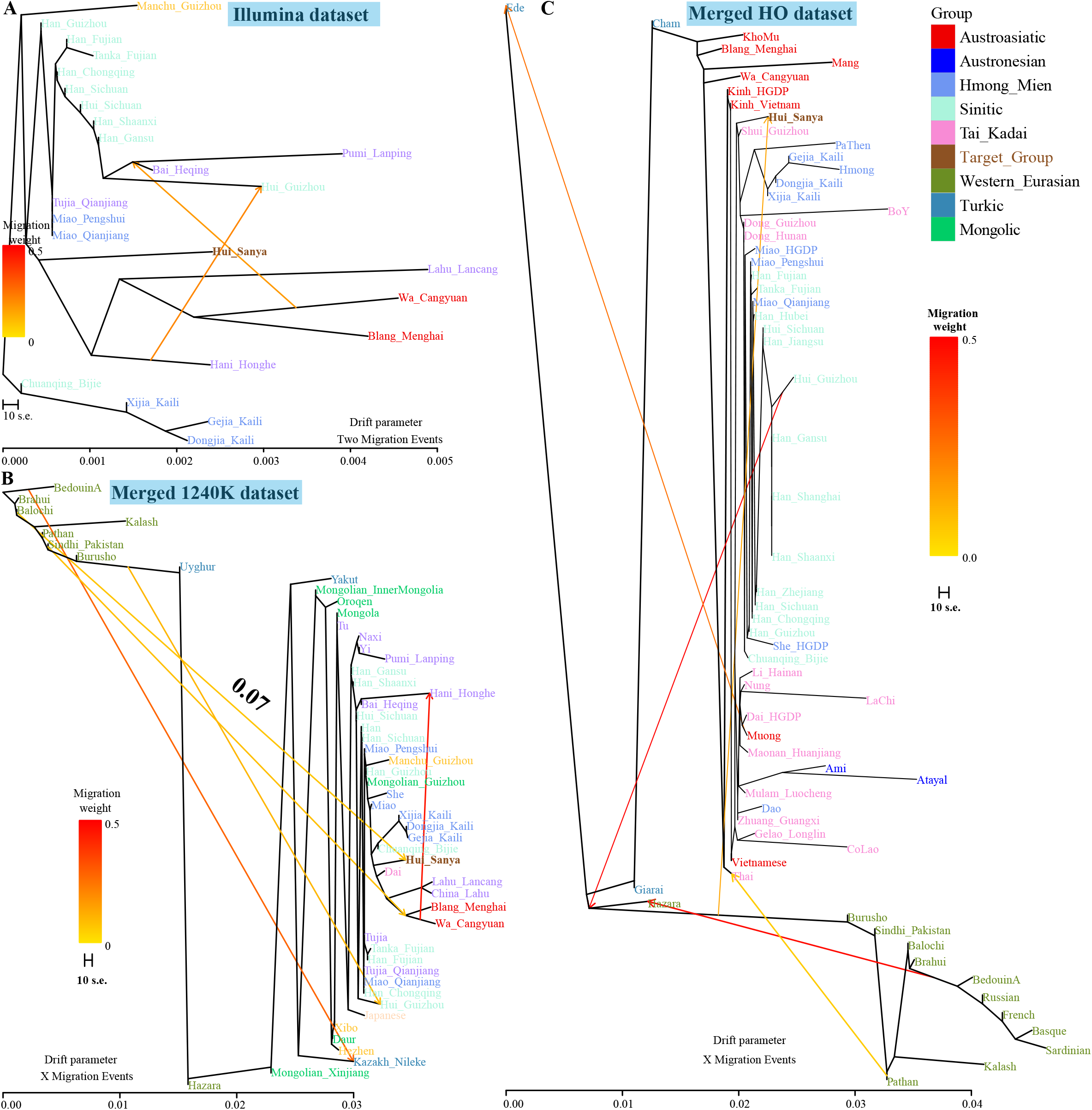
Phylogenetic relationship reconstruction based on the maximum likelihood and allele frequency distribution. (**A**), Population splits and two gene flow events among 24 Chinese populations based on the merged 1240K dataset. (**B**), Populations splits and five admixture events among 54 central and eastern Eurasian populations based on the merged 1240K dataset. (**C**), TreeMix-based phylogenetic tree with six migration events among 64 populations based on the merged HO dataset.

To directly explore the ancestral source and corresponding admixture proportion of SYH people, we modeled their gene pool via the three-way admixture qpAdm models (**Figure 5A**). In our findings based on the *f_4_*-statistics and TreeMix-based phylogenetic topology, we also found a close genetic relationship between SYH and Austroasiatic people in Vietnam, thus, we used Cham people (green ancestry) as one of three sources, the other two sources from western Eurasian and South China, we found that SYH people were the mixture results of major ancestry related to southern Chinese populations, including Hmong-Mien, Tai-Kadai and Han Chinese people, and minor ancestry related to Cham and western Eurasians. We provided direct genetic evidence supporting three ancestral sources contributed to the formation of modern SYH people.

**Figure 5.**
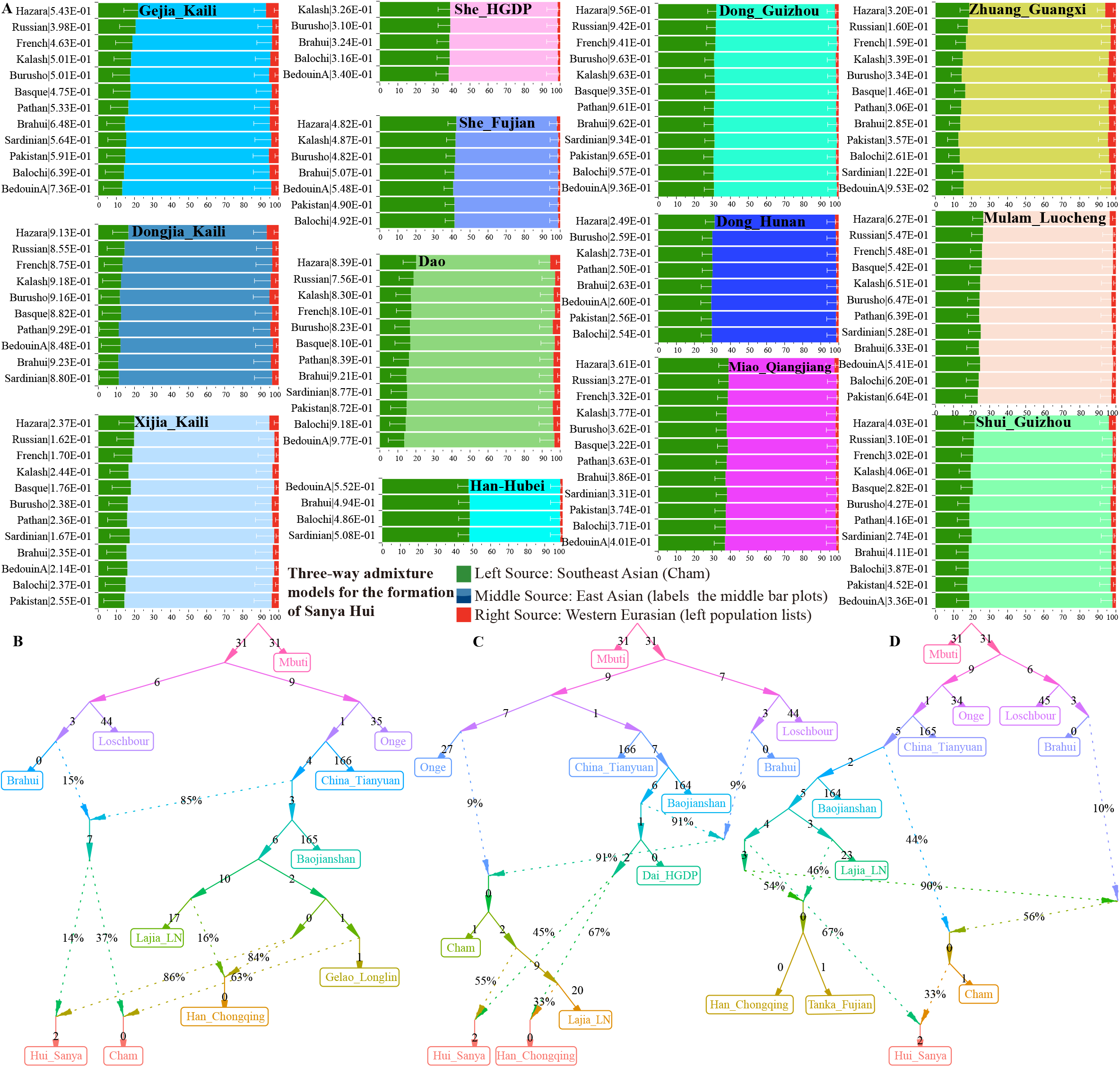
Demographic admixture models for Sanya Hui people. (**A**), Three-way admixture models for Sanya Hui people with different sources of Southeast Asian Cham (first green ancestry), western Eurasians (third ancestry labeled in the left population lists) and East Asians (middle ancestry marked in each group). (**B~D**), Deep population admixture models of Sanya Hui and other related modern and ancient populations reconstructed via qpGraph. Dot line denoted the admixture events, and the number was the genetic drift (1000 times of original values) and the percent number was denoted as the admixture proportion.

We also formally explored the deep population history of the Hui people and their relatives via qpGraph models with population divergence and admixture among modern and ancient populations. Firstly, we parallelly modeled the formation of Hui and Cham people (**Figure 5B**), we found that Hui and Cham people could be modeled as a mixture of western Eurasian (Brahui: 0.021 and 0.056), deep early Asian (0.119 and 0.315) and southern Chinese ancestry related to Gelaos (0.86 and 0.63). We also found that Han Chinese populations from Chongqing could be well-fitted by the Gelao-Lajia model (Gelao_Longlin: 0.84; Lajia_LN: 0.16). When modeled Cham as the ancestor of the Hui people (**Figure 5C**), we found that SYH could be modeled as 0.55 ancestry related to Cham and additional ancestry from Chinese Dai people. We also modeled Chongqing Han in this model, which could be fitted as 0.33 ancestry from Lajia and the remaining from Dai. Furthermore, to confirm the stability of the aforementioned models (**Figure 5D**), we built the third model, which also provided evidence for the derived ancestry of SYH people from Cham and southern Chinese populations. We should be cautious with the estimated admixture proportions in these three models, which at least consistently showed that modern SYH people derived ancestry from populations related to Cham and southern Chinese populations, the western affinity of Hui people might be indirectly introduced via Cham people, but we cannot confirm the possibility of additional western admixture into the descendants of Cham, as historic documents showed that Hui people migrated from North China also participated in the formation of SYH people.

### Sex-biased admixture

We assessed the potentially existing sex-biased admixture in the formation of SYH people via three ways (derived ancestry from autosomal, Y-chromosomal and mtDNA evidence). We successfully fitted ten two-way admixture models with Thai as the southern source and others from EA as the northern source, in which the estimated ancestry contribution from Thai was significantly higher based on the autosomes than based on the X-chromosome (**Figures 6A~B**). Additional well-fitted two-way admixture models with Malay as the southern source also confirmed the higher autosome-based admixture proportion in SYH than X-chromosome-based proportion, which supported the male-dominant population admixture from SEA. We then compared the ancestry frequency distribution between southern (Sanya), central (Sichuan and Guizhou) Hui and Han people (**Figures 6C~D**), uniparental landscapes in these populations showed that maternal F haplogroup and paternal O1a and O1b were dominant lineages in SYH. Results from the Chi-squared test based on the allele frequency of uniparental lineages showed that maternal and paternal genetic contribution in ethnically different Han and Hui people, and geographically different Hui people were significantly diverse. Different from the genetic history of Guizhou Hui and Han people, SYH people harbored more paternal ancestry from CSA, such as western Eurasian-related J2b2a2b and C1b1a paternal lineages frequently observed in SYH. These identified sex-specific contributions from paternal and maternal legacy demonstrated that the general admixture process of SYH was sex-biased with an excess of Southeast Asian indigenous males and East Asian females. Previous studies based on whole-genome sequencing, genome-wide SNPs, Y-chromosomal and mitochondrial loci also have shown the sex-based admixture process in Hui people (Liu et al., 2021b; Ma et al., 2021; Wang et al., 2020a). It is interesting to find that the genetic contribution from Cham people to SYH was mediated via females.

**Figure 6.**
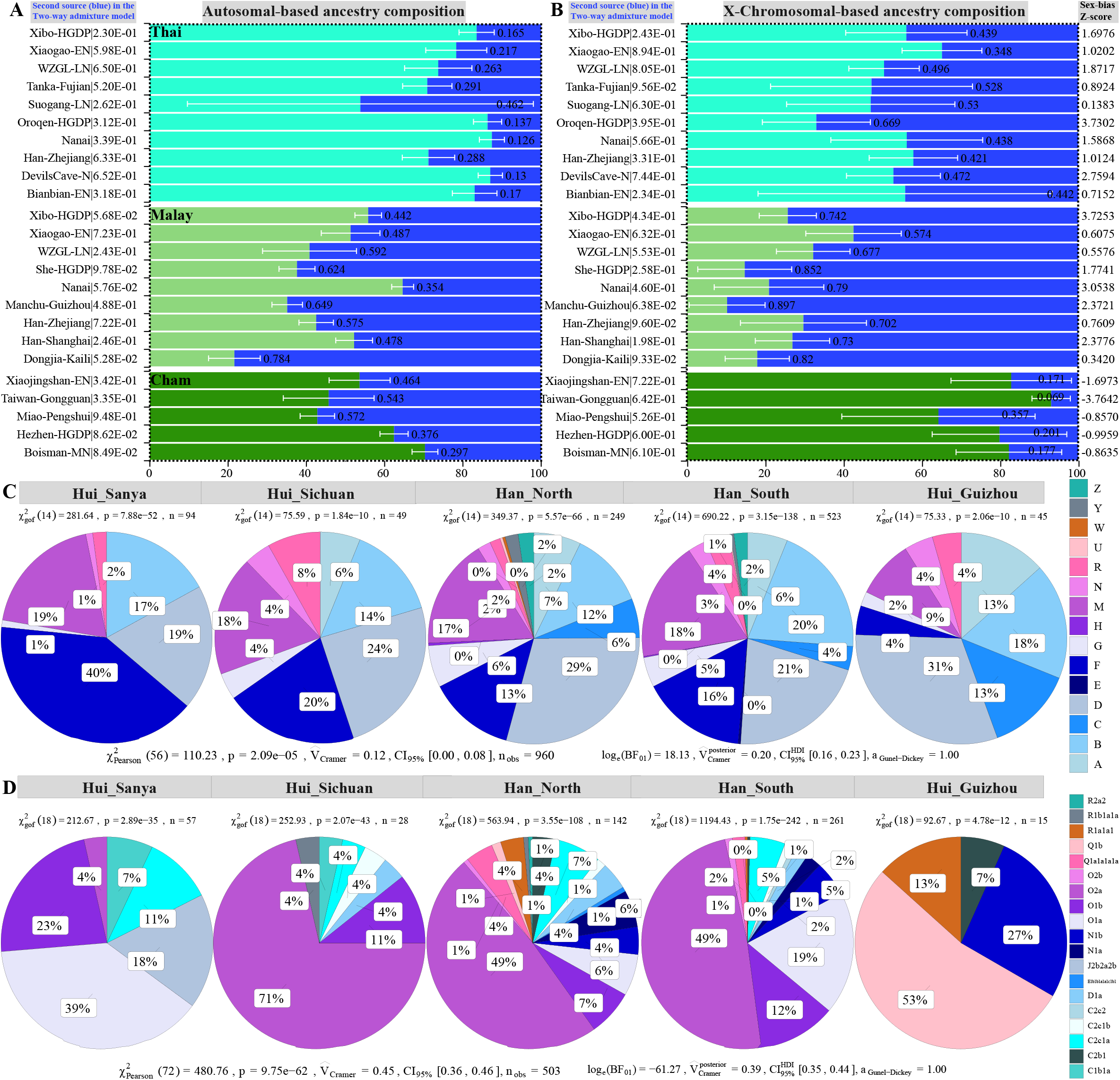
Evidence from autosomal and uniparental chromosomes showed the sex-biased admixture process. (**A**), Two-way admixture models with Thai, Malay, and Cham as the Southeast Asian ancestral source of SYH people and other East Asians as the second source based on the autosome (1~22). (**B**), Two-way admixture models showed the admixture ancestry proportion based on the X-chromosome. The left number was the p_values of the rank1. The right number was the Z-scores which showed the differences between the autosomal and X-chromosomal ancestry proportion, in which the positive values showed that more southeast males participated in the formation of SYH and negative values showed more East Asian males participated in the formation. (**C~D**), Similarities and differences of the frequency distribution of the mtDNA-haplogroups (**C**) and Y-haplogroups (**D**) of Hui and northern and southern Han Chinese populations.

### Haplotype-based finer-scale population genetic structure

Estimated pairwise IBD fragments showed a close genetic affinity between SYH and reference Huis, followed by Cham, Giarai, and southern Chinese populations (She, Tanka, Dai, and Maonan). When we used other populations from EA and CSA as the ancestral donors to paint the chromosome of SYH people (**Figure 7A**), we found that Han Chinese from South China and geographically close Tai-Kadai people contributed more DNA fragments to SYH people. Followingly, we used SYH as the ancestral donor to paint all other included phased population’s chromosomes from Southeast Asians and East Asians and we found that Tai-Kadai-speaking Dai and southern Han Chinese received the longest DNA chunks donated by SYH people. We also explored the finer-scale population structure via fine STRUCTURE based on the two different datasets: the merged HO and the merged 1240k datasets among East Asians and Central Asians (**Figures 7B~D**), we identified one separated cluster branch of SYH, which was localized between western and eastern Eurasian populations.

**Figure 7.**
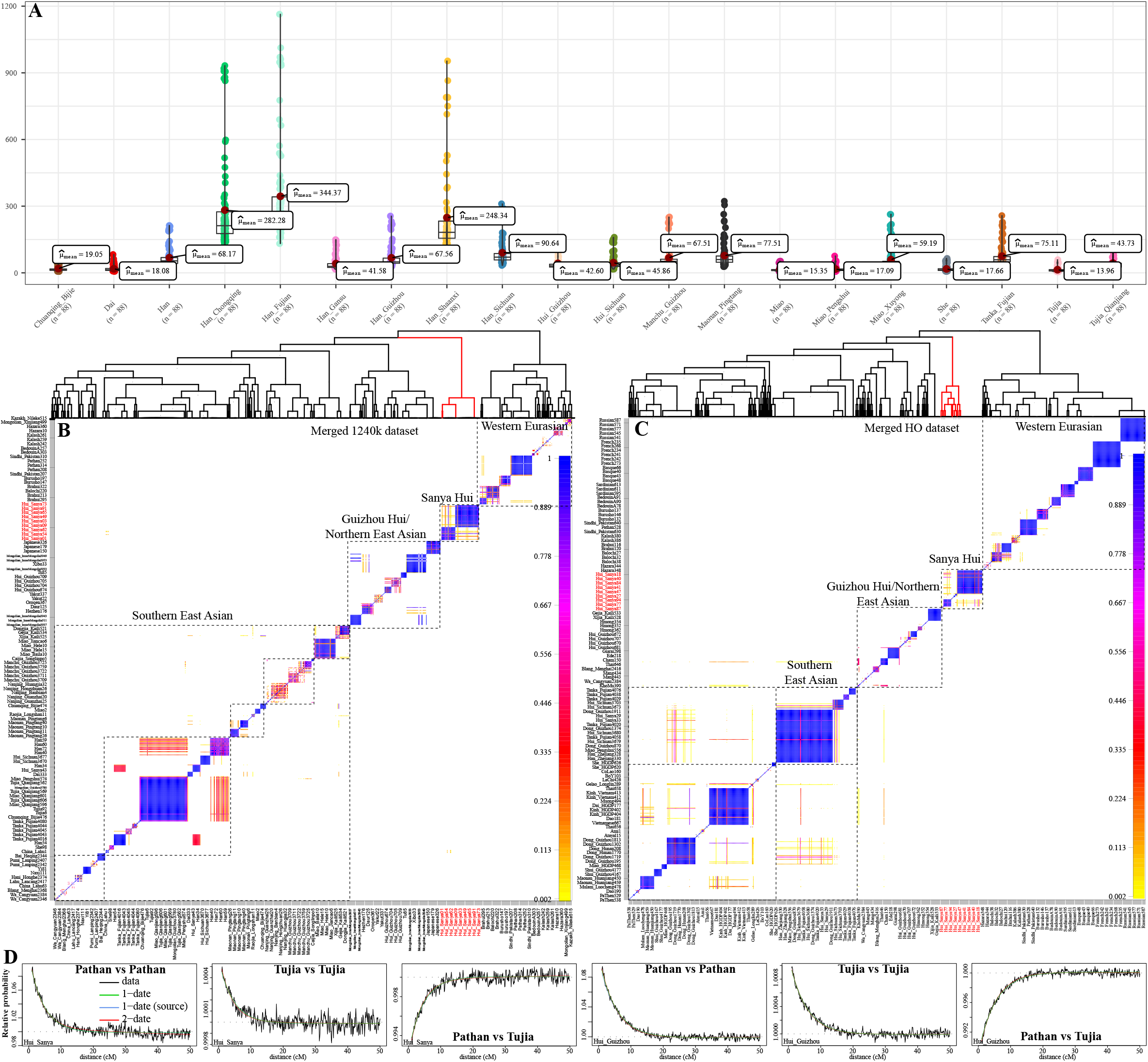
Finer-scale population structure inferred from the dense haplotype data. (**A**), ChromoPainter results showed that SYH people copied ancestral chunk length from other East Asians. (**B~C**), Pairwise coincidence matrixes based on the merged 1240K and HO datasets showed the unique genetic structure of SYH people. (**D**), The inferred coancestry curves showed the relative probability that two IBD chunks at a certain genetic distance (X-axis) were copied from a pair of surrogate Pathan and Tujia after removing Han Chinese populations in the GLOBETROTTER analysis.

To explore the admixture events based on the haplotype data, we conducted GLOBETTRTER analysis based on the sharing haplotypes. Among the merged 1240K dataset, we found that the one-date-one-way admixture was the best-fitted model (fit. quality. 1 event: 0.9986). SYH people were the mixed results of minor ancestral source (0.05) related to Sindhi Pakistan and major ancestral source related to Han at the 28 years ago. We used the same strategy to date and describe the admixture processes of the Guizhou and Sichuan Hui people. We identified the best-fitted one-date-one-way model for Guizhou Hui people with the minor western source related to Pathan (0.07) and major eastern source related to Sichuan Han (0.93) at 20 years ago. Sichuan Hui people showed the same admixture source and admixture time as Sichuan Hui, which were the mixture results via minor Pathan source (0.03) and major East Asian source (0.97) at 22 years ago. We also obtained consistent admixture times based on the ALDER methods (**Table S9**). After we removed Han Chinese from the donor populations, we identified robust and consistent decay curves of the two IBD fragments at a certain genetic distance originated from the common ancestor (**Figure 7D**), which showed that other CSA populations rather than Pathan and other East Asian rather than Han (Tujia) people could be also modeled as the surrogates of Hui’s ancestors.

Finally, we explored the natural selection signatures of SYH people and identified the mechanisms of biological adaptation of the island environments. We used the top twenty-one genes included at least three functional SNPs with natural selection signatures to conduct Functional Annotation Clustering in the Database for Annotation, Visualization and Integrated Discovery (DAVID) (**Figure 8A**). We identified three clusters based on these top-selected genes respectively associated membrane function (Enrichment Score: 0.34: Transport, membrane, an integral component of membrane/plasma membrane, Transmembrane/Transmembrane helix, topological domain: Cytoplasmic, transmembrane region and Membrane), Zinc-related function (0.32: zinc ion binding, Zinc-finger, Zinc and Metal-binding) and extracellular complex functions ( including extracellular exosome, Signal/Signal peptide, plasma/cell membrane, Glycoprotein/glycosylation site). Among the top-identified genes, *FERM domain-containing 4A* (FRMD4A) codes proteins of cytoplasm, cytoskeleton, bicellular tight junction, which is responsible for protein binding and bridging and associated with the biological process of establishment of epithelial cell polarity. MUC22 gene was the top natural-selected candidate located in chromosome 6 and encodes protein associated with the integral component of the membrane. *Tripartite motif-containing 31* (TRIM31) is also localized in chromosome 6 and encodes intracellular, mitochondrion, cytosol proteins responsible for protein binding, zinc ion binding, and ligase activity. *Family with sequence similarity 179 member A* (FAM179A) and *Ring finger protein 39* (RNF39) are also localized in chromosome 6 that associated with the molecular function of zinc ion binding in the cytoplasm. We further calculated the iHS scores of Hui people and XPEHH values compared with southern Han Chinese. We also found the natural-selected signatures in the *Human leukocyte antigens* (HLA) and *Major Histocompatibility Complex* (MHC)-related genes, which are associated with the immune functions and rapidly adapt to new environments. Although we identified western Eurasian mixed ancestry in SYH people, we did not observe the natural-selected signatures from the LCT and pigmentation-related genes predominant in Western Eurasians. The SYH people shared similar general patterns of natural-selected pattens with eastern Eurasians. Furthermore, we calculated the other XPEHH values in Hui people referenced to southern Hans and also assessed their iHS scores and found that more consistent natural-selected signatures were identified based on two sets of the XPEHH-based candidates (**Figure 8B**) rather than the shared signals between XPEHH-based candidate genes and iHS-based candidate genes. We further conducted enrichment analysis based on the three sets of loci candidates and explored the involved biological pathways, gene and gene interactions, and the proprotein interactions (**Figures 8C~G**). We found these identified loci associated with fourteen biological processes (regulation of cell morphogenesis (GO: 0022604), circulatory system process (GO: 0003013), positive regulation of cell-substrate adhesion (GO: 0010811), antigen processing and presentation of endogenous peptide antigen via MHC class I (GO: 0019885), regulation of plasma membrane-bounded cell projection organization (GO: 0120035), negative regulation of cellular component organization (GO: 0051129), female pregnancy (GO: 0007565), eye photoreceptor cell differentiation (GO: 0001754), calcium ion transmembrane transport (GO: 0070588), neuron recognition (GO: 0008038), learning (GO: 0007612), Notch signaling pathway (GO: 0007219), filopodium assembly (GO: 0046847) and cell fate determination (GO: 0001709)). Three KEGG pathways (ABC transporters (hsa02010), Drug metabolism - cytochrome P450 (hsa00982), AGE-RAGE signaling pathway in diabetic complications (ko04933)) and three reactome gene sets (Generation of second messenger molecules (R-HSA-202433), Transport of small molecules (R-HSA-382551), Collagen degradation (R-HSA-1442490)) were also associated with our identified natural-selected loci. Our results suggested that SYH people underwent a similar process of natural selection with other East Asians, such as Han Chinese populations.

**Figure 8.**
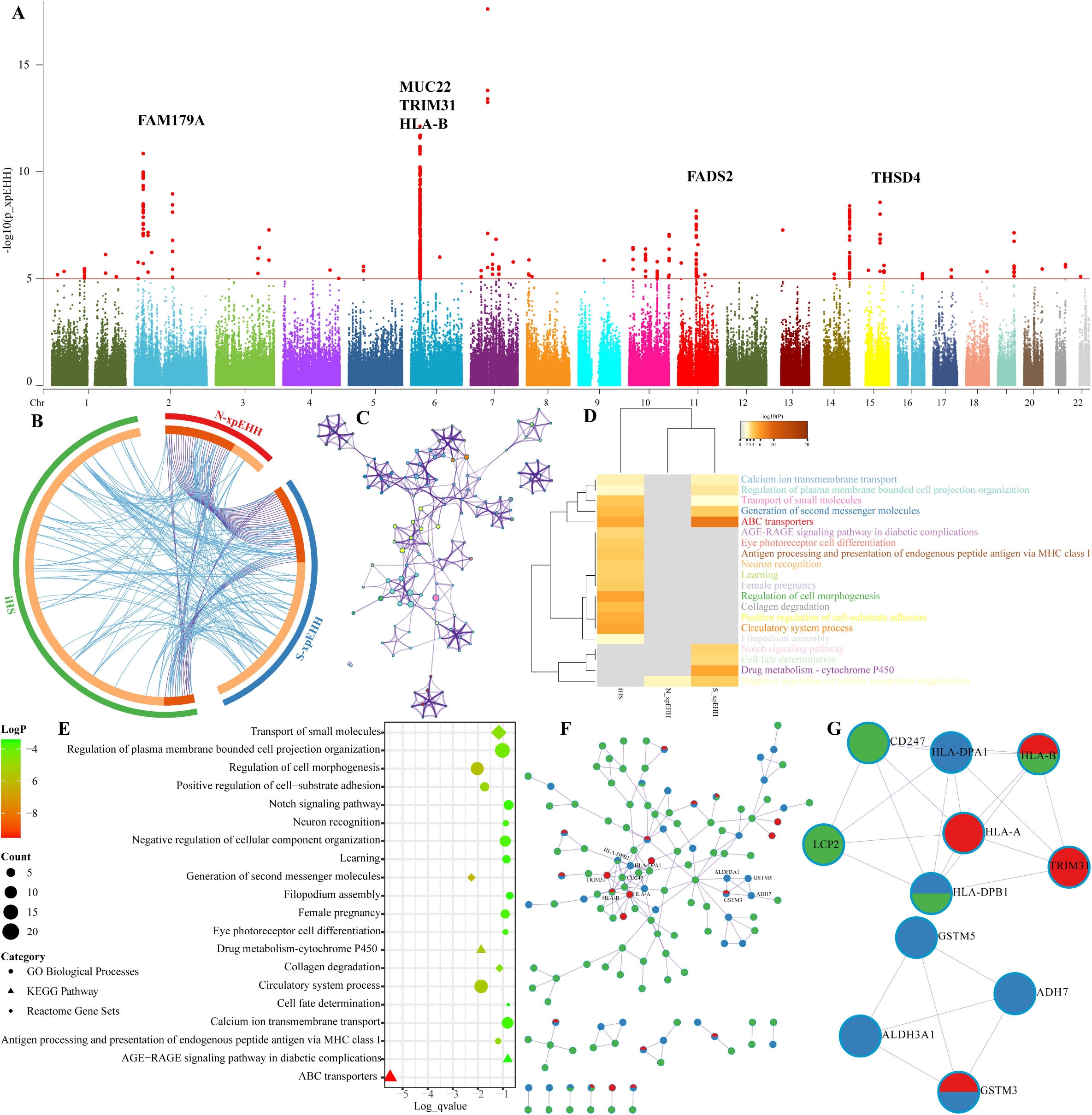
Natural-selected signatures of Sanya Hui people. (**A**), Manhattan plots showed the p values of the XPEHH. (**B**), Overlaps among the included and shared gene lists, where blue curves link genes that belong to the same enriched ontology term. The inner-circle represents gene lists, where hits are arranged along the arc. Genes that hit multiple lists are colored in dark orange, and genes unique to a list are shown in light orange. (**C**), Network of enriched terms colored by cluster-ID. (**D**), Heatmap of enriched terms across three sets of input gene lists focused on the Hui people, colored by p-values. (**E**), Top 20 clusters with their representative enriched terms. (**F~G**), Protein-protein Interaction Enrichment Analysis.

## Discussion

Hui was the third largest ethnic group in China followed behind the Han and Zhuang people. The fine-scale genetic structure, demographic models of Hui people and their regional biological adaptation were important for the study design in the phenome-wide association studies (PheWASs) and genome-wide association studies (GWASs). Comparisons with the comprehensive genetic studies focused on the genetic diversity of Han Chinese populations (Chen et al., 2009; Chiang et al., 2018; GuangūLin He et al., 2021; He et al., 2021; Liu et al., 2018; Xu et al., 2009; Yao et al., 2021), works on inferring Hui’s demographic history and surveying their genetic variations keep in its infancy. However, to provide a complete genomic landscape of representative ethnolinguistically and geographically diverse populations in the Chinese Population Genomic Diversity Project (CPGDP) and corresponding Chinese Trans-omics for Precision Medicine (C-TOPMed) via whole-genome sequencing, it is important to investigate the general but complete patterns of East Asian’s genetic diversity and structure via array-based scanning (Cavalli-Sforza, 1998). Here, we reported the first batch of genome-wide SNP data from the autosome, X/Y-chromosome, and mtDNA data of SYH people and presented the comprehensive population genetic analyses to characterize their admixture history and biological adaptation. We not only identified a significant genetic differentiation between SYH and central Chinese Hui people from Guizhou and Sichuan but also found the differentiated genetic admixture history between SYH people and indigenous Hainan Tai-Kadai populations and Vietnam Cham people. Our findings reported here suggested the more complex landscape of the demographic history of Chinese Hui people than previously reported two-source admixture models (Liu et al., 2021b; Ma et al., 2021; Wang et al., 2020a) and provided insights into the formation of southernmost Hui people, which was formed via a mixture of tripartite sources from EA, CV and CSA.

Compared with Eurasian populations via pairwise Fst genetic distances and outgroup-*f_3_*-statistics values, SYH people harbored a close genetic relationship with Southeast Asian Austroasiatic-speaking Kinh and Vietnamese, and also with geographically close Tai-Kadai people. These identified patterns were further confirmed via the clustering patterns between SYH and their neighbors revealed in the PCA and ADMIXTURE. We consistently obtained the minor ancestry sources from Pakistani populations in CSA based on the GLOBETROTTER analysis focused on Sanya, Guizhou, and Sichuan Hui populations. Besides, we also identified the western Eurasian affinity compared with geographically close populations (Han or Li) in the *f_4_*-statistics and qpAdm-based analysis, which was consistent with previously reported western Eurasian admixture signatures in the central Chinese Hui people. For example, statistically negative *f_4_*((Western Eurasians, Eastern Eurasians; SYH, Mbuti) quantitatively illuminated SYH people shared more alleles with East Asians and Southeast Asians compared with Western Eurasians, which was consistent with the higher estimated Eastern Eurasian-related ancestry proportion in the qpAdm and qpGraph models. Recent genetic studies based on the central Guizhou and Sichuan Huis, northwestern Ningxia, and Xinjiang Huis also showed eastern Eurasian affinity of their studied Hui people (Liu et al., 2021b; Ma et al., 2021; Wang et al., 2020a). Taken together, we concluded that geographically different Chinese Hui people derived their primary ancestry from East Asians although they remain unique Islam culture and dietary habits. Our reported genetic structure supported that the key role of the cultural diffusion model and limited demic diffusion model influenced the formation of the Chinese Hui people.

Although the reported genetic structure among Hui people from Sanya, Guizhou, and Sichuan reflected the common origin of western ancestry sources related to Central Asian populations and East Asian affinity, we also identified the differentiated influence of gene influxes from geographically different indigenous people into the Hui people in descriptive analyses (PCA, ADMIXTURE and pairwise Fst values). Guizhou Hui was reported as a unique cline in the East Asian-scale PCA, and Sichuan and Gansu Huis clustered closely with northwestern Chinese populations (Liu et al., 2021b; Ma et al., 2021; Wang et al., 2020a). However, SYH people formed a separated cluster in the PCA, a unique ancestral component in the model-based ADMIXTURE and possessed a close relationship with Vietnam populations in the Fst and outgroup-*/3*-based affinity indexes, indicating that the assimilation with different East Asian ancestry sources influenced the genetic structure of Hui people. Indeed, genetic analyses focused on Hui people from Xinjiang, Ningxia, Sichuan, and Guizhou had demonstrated that Hui people from central and northwestern China possessed a strong genetic affinity with northern East Asians (Liu et al., 2021b; Ma et al., 2021; Wang et al., 2020a). Previous genetic analyses focused on modern ethnolinguistically diverse populations not only identified the genetic distinction between northern and southern East Asians but also observed at least five genetic clines among Chinese populations (Gao et al., 2020; Wang, C.C. et al., 2021), which included northern Tibeto-Burman cline among Tibetan Plateau populations, Tungusic and Mongolic cline among Amur River populations, Hmong-Mien cline among southwestern populations, Austronesian cline among Taiwan indigenous populations and north-to-south cline consisting of Han Chinese populations and Tai-Kadai speakers. Recent ancient genomes also revealed the differentiated ancestries among these geographical regions, at least including Guangxi ancestry related to Longlin, Baojianshan and Dushan (Wang, T. et al., 2021), Fujian ancestry related to early Neolithic Qihe and latter Neolithic Tanshishan and Xitoucun (Yang et al., 2020), Shandong ancestry related to Xiaogao, Xiaojingshan and Bianbian (Yang et al., 2020), Henan and Gansu ancestry related to Yangshao and Longshan farmers (Ning et al., 2020), Tibetan ancestry related to Chokhopani (Jeong et al., 2016), and Amur River ancestry related to AR14K (Wang, T. et al., 2021). Obvious, genetic analyses showed the observed landscape of southernmost Chinese Hui were mixed of one western Eurasian ancestry and one eastern southern East Asian ancestry and one Southeast Asian ancestry. Generally, modern Hui people were formed via the minor western Eurasian ancestry who migrated to China since the Tang dynasty and then assimilated with geographically different East Asian indigenous populations, which was obtained based on the estimated ALDER and GLOBETROTTER-based admixture times and historic documents.

Beyond the reported general origin and differentiated admixture history of the Hui people, our study provided robust genetic evidence to resolve the controversial opinion that SYH people were the direct descendants of Cham people (Li et al., 201). Our results showed that modern Cham people had a distinct relationship with SYH in the PCA and ADMIXTURE results. The observed large pairwise Fst genetic distance between SYH and Cham and the smallest outgroup-*f_3_* values further confirmed their distant genetic relationship. QpAdm-based three-way admixture models with Cham as one of the sources and qpGraph-based models further showed the limited gene flow from Cham’s gene pool, and SYH people obtained excessive additional ancestry from East Asians. Thus, our population genetic analysis supported that modern SYH people were formed via three ancestral sources from CSA, SEA and EA and rejected that the SYH people were the direct descendants of the ancient Cham people. Our reported patterns of natural-selected signals in SYH people showed similar patterns as the East Asians, which further confirmed the strong genetic influence of East Asians on modern SYH people.

## Conclusion

Historic documents suggested that modern Hui people were assimilated by Islam and might be the descendants of ancient Silk Road immigrants who migrated into China from CSA, the Middle East, etc. To provide the finer-scale genetic structure of Hainan Hui and reconstruct their demographic history, we genotyped genome-wide SNPs in 94 Hainan Hui people. We identified one unique genetic structure of SYH people, which was different from Vietnam Cham and geographically different Hui people from Guizhou and Sichuan. SYH people were modeled as the mixed population with three ancestral sources respectively from CSA, SEA, and EA. In comparison with modern and ancient populations from eastern Eurasia, we found the genetic makeup of Chinese Hui populations has been strongly influenced by the assimilation with geographically different East Asian indigenous sources. Our GLOBETTRETOR-based admixture times showed that massive genetic admixture between eastern and western sources occurred during the Tang dynasty and the Five Dynasties and Ten Kingdoms period, which should be further confirmed via whole-genome sequencing data from SYH people. Natural-selected signals from genes associated with membrane functions and immune response promoted the biological adaptation of SYH people. Finally, we identified the sex-biased admixture process based on the qpAdm-based autosomal signals, Y-chromo somal, and mitochondrial lineage s. Taken together, tripartite ancestral sources contributed genetic materials to modern SYH people, which promoted them to rapidly adapt to the hot tropical environments.

## Acknowledgments

This work was funded by the Science and Technology Program of Guangzhou, China (2019030016), the Project funded by China Postdoctoral Science Foundation (2021M691879), Social Science Foundation Project of Fujian Province (FJ2020B131), National Social Science Foundation of China (20&ZD248), the “Double First Class University Plan” key construction project of Xiamen University (the origin and evolution of East Asian populations and the spread of Chinese civilization), National Natural Science Foundation of China (NSFC 31801040), Nanqiang Outstanding Young Talents Program of Xiamen University (X2123302), the Major project of National Social Science Foundation of China (20&ZD248), the European Research Council (ERC) grant to Dan Xu (ERC-2019-ADG-883700-TRAM). S. Fang and Z. Xu from the Information and Network Center of Xiamen University are acknowledged for their help with high-performance computing. We thank Prof. Wibhu Kutanan in Khon Kaen University, Prof. Mark Stoneking and Dr. Dang Liu in Max Planck Institute for Evolutionary Anthropology for sharing genome-wide SNP data from Vietnam, Thailand, and Laos.

## Legends of Supplementary Tables

Table S1. Pairwise Fst genetic distances among 24 Chinese populations based on the merged Illumina dataset.

Table S2. Pairwise Fst genetic distances among 204 Eurasian populations based on the merged HO dataset.

Table S3. Pairwise outgroup-*f_3_* values among 250 Eurasian populations were estimated via *f_3_*(Source1. Source2; Mbuti).

Table S4. Admixture signatures inferred from the admixture-*f_3_*-statistics in the form *f_3_*(Source1, Source2; Sanya Hui).

Table S5. The asymmetric relationship between Sanya Hui and other Hui people from Guizhou and Sichuan was inferred via *f_4_*(Hui1, Hui2; reference populations, Mbuti).

Table S6. Results of *f_4_*-statistics in the form *f_4_*(Refrence1, Reference2; Sanya Hui, Mbuti) showed the East Asian affinity of Sanya Hui people.

Table S7. Results of *f_4_*-statistics in the form *f_4_*(Refrencel, Sanya Hui; Reference2, Mbuti) showed the genetic relationship between Sanya Hui and their predefined surrogates of ancestral sources based on the linguistic and historic documents.

Table S8. Pairwise p values in the pairwise qpWave analysis among 129 populations.

Table S9. ALDER results showed the admixture times with the predefined ancestral population pairs.

## Reference

Alexander, D.H., Novembre, J., Lange, K., 2009. Fast model-based estimation of ancestry in unrelated individuals. Genome Res 19(9), 1655–1664. https://doi.org/10.1101/gr.094052.109.

Andaya, L.Y., 2008. Leaves of the same tree. University of Hawaii Press.

Browning, B.L., Browning, S.R., 2011. A fast, powerful method for detecting identity by descent. Am J Hum Genet 88(2), 173–182. https://doi.org/10.1016/j.ajhg.2011.01.010.

Browning, B.L., Browning, S.R., 2013. Improving the accuracy and efficiency of identity-by-descent detection in population data. Genetics 194(2), 459–471. https://doi.org/10.1534/genetics.113.150029.

Cavalli-Sforza, L.L., 1998. The Chinese human genome diversity project. Proc Natl Acad Sci U S A 95(20), 11501–11503. https://doi.org/10.1073/pnas.95.20.11501.

Chang, C.C., Chow, C.C., Tellier, L.C., Vattikuti, S., Purcell, S.M., Lee, J.J., 2015. Second-generation PLINK: rising to the challenge of larger and richer datasets. Gigascience 4, 7. https://doi.org/10.1186/s13742-015-0047-8.

Chen, C., Li, Y., Tao, R., Jin, X., Guo, Y., Cui, W., Chen, A., Yang, Y., Zhang, X., Zhang, J., Li, C., Zhu, B., 2020. The Genetic Structure of Chinese Hui Ethnic Group Revealed by Complete Mitochondrial Genome Analyses Using Massively Parallel Sequencing. Genes (Basel) 11(11). https://doi.org/10.3390/genes11111352.

Chen, J., He, G., Ren, Z., Wang, Q., Liu, Y., Zhang, H., Yang, M., Zhang, H., Ji, J., Zhao, J., Guo, J., Zhu, K., Yang, X., Wang, R., Ma, H., Wang, C.C., Huang, J., 2021. Genomic Insights Into the Admixture History of Mongolic- and Tungusic-Speaking Populations From Southwestern East Asia. Front Genet 12(880), 685285. https://doi.org/10.3389/fgene.2021.685285.

Chen, J., Zheng, H., Bei, J.X., Sun, L., Jia, W.H., Li, T., Zhang, F., Seielstad, M., Zeng, Y.X., Zhang, X., Liu, J., 2009. Genetic structure of the Han Chinese population revealed by genome-wide SNP variation. Am J Hum Genet 85(6), 775–785. https://doi.org/10.1016/j.ajhg.2009.10.016.

Chen, L., 1999. Zhongguo minzushi gangyao (The compendium of Chinese nationality histories). China Financial & Economic Press, Beijing, China.

Chiang, C.W.K., Mangul, S., Robles, C., Sankararaman, S., 2018. A Comprehensive Map of Genetic Variation in the World’s Largest Ethnic Group-Han Chinese. Mol Biol Evol 35(11), 2736–2750. https://doi.org/10.1093/molbev/msy170.

Deng, Y-j., Zhu, B.-f., Shen, C.-m., Wang, H.-d., Huang, J.-f., Li, Y.-z., Qin, H.-x., Mu, H.-f., Su, J., Wu, J., 2011. Genetic polymorphism analysis of 15 STR loci in Chinese Hui ethnic group residing in Qinghai province of China. Molecular biology reports 38(4), 2315–2322.

Dillon, M., 2013. China’s Muslim Hui community: migration, settlement and sects. Routledge.

Edmondson, J.A., Gregerson, K.J., 1993. Tonality in Austronesian languages. University of Hawaii Press.

Elhaik, E., Tatarinova, T., Chebotarev, D., Piras, I.S., Maria Calo, C., De Montis, A., Atzori, M., Marini, M., Tofanelli, S., Francalacci, P., Pagani, L., Tyler-Smith, C., Xue, Y., Cucca, F., Schurr, T.G., Gaieski, J.B., Melendez, C., Vilar, M.G., Owings, A.C., Gomez, R., Fujita, R., Santos, F.R., Comas, D., Balanovsky, O., Balanovska, E., Zalloua, P., Soodyall, H., Pitchappan, R., Ganeshprasad, A., Hammer, M., Matisoo-Smith, L., Wells, R.S., Genographic, C., 2014. Geographic population structure analysis of worldwide human populations infers their biogeographical origins. Nat Commun 5, 3513. https://doi.org/10.1038/ncomms4513.

Gao, Y., Zhang, C., Yuan, L., Ling, Y., Wang, X., Liu, C., Pan, Y., Zhang, X., Ma, X., Wang, Y., Lu, Y., Yuan, K., Ye, W., Qian, J., Chang, H., Cao, R., Yang, X., Ma, L., Ju, Y., Dai, L., Tang, Y., Han, K.I., Zhang, G., Xu, S., 2020. PGG.Han: the Han Chinese genome database and analysis platform. Nucleic Acids Res 48(D1), D971–D976. https://doi.org/10.1093/nar/gkz829.

Gautier, M., Klassmann, A., Vitalis, R., 2017. rehh 2.0: a reimplementation of the R package rehh to detect positive selection from haplotype structure. Molecular ecology resources 17(1), 78–90. https://doi.org/10.1111/1755-0998.12634.

Gladney, D.C., 1996. Muslim Chinese: ethnic nationalism in the People’s Republic. Harvard Univ Asia Center.

Gladney, D.C., 1998. Ethnic identity in China: the making of a Muslim minority nationality. Harcourt Brace College Publishers Fort Worth, TX.

Grant, A., Sidwell, P., 2005. Chamic and beyond: studies in mainland Austronesian languages. Pacific Linguistics, Research School of Pacific and Asian Studies, The ….

GuangūLin He, Li, Y.X., Wang, M.G., Zou, X., Yeh, H.Y., Yang, X.M., Wang, Z., Tang, R.K., Zhu, S.M., Guo, J.X., Luo, T., Zhao, J., Sun, J., Xia, Z.Y., Fan, H.L., Hu, R., Wei, L.H., Chen, G., Hou, Y.P., ChuanūChao, W., 2021. Fineū scale genetic structure of Tujia and central Han Chinese revealing massive genetic admixture under language borrowing. Journal of Systematics and Evolution 59(1), 1–20.

Guo, F., 2017. Population genetics for 17 Y-STR loci in Hui ethnic minority from Liaoning Province, Northeast China. Forensic Sci Int Genet 28, e36–e37. https://doi.org/10.1016/j.fsigen.2017.02.011.

He, G., Wang, Z., Wang, M., Luo, T., Liu, J., Zhou, Y., Gao, B., Hou, Y., 2018. Forensic ancestry analysis in two Chinese minority populations using massively parallel sequencing of 165 ancestry-informative SNPs. Electrophoresis 39(21), 2732–2742. https://doi.org/10.1002/elps.201800019.

He, G.L., Wang, M.G., Li, Y.X., Zou, X., Yeh, H.Y., Tang, R.K., Yang, X.M., Wang, Z., Guo, J.X., Luo, T., Zhao, J., Sun, J., Hu, R., Wei, L.H., Chen, G., Hou, Y.P., Wang, C.C., 2021. Fineūscale northūtoūsouth genetic admixture profile in Shaanxi Han Chinese revealed by genome wide demographic history reconstruction. Journal of Systematics and Evolution, 0–. https://doi.org/10.1111/jse.12715.

Hellenthal, G., Busby, G.B.J., Band, G., Wilson, J.F., Capelli, C., Falush, D., Myers, S., 2014. A genetic atlas of human admixture history. Science 343(6172), 747–751. https://doi.org/10.1126/science.1243518.

Jeong, C., Ozga, A.T., Witonsky, D.B., Malmstrom, H., Edlund, H., Hofman, C.A., Hagan, R.W., Jakobsson, M., Lewis, C.M., Aldenderfer, M.S., Di Rienzo, A., Warinner, C., 2016. Long-term genetic stability and a high-altitude East Asian origin for the peoples of the high valleys of the Himalayan arc. Proc Natl Acad Sci U S A 113(27), 7485–7490. https://doi.org/10.1073/pnas.1520844113.

Lawson, D.J., Hellenthal, G., Myers, S., Falush, D., 2012. Inference of population structure using dense haplotype data. PLoS Genet 8(1), e1002453. https://doi.org/10.1371/journal.pgen.1002453.

Li, D.N., Wang, C.C., Yang, K., Qin, Z.D., Lu, Y., Lin, X.J., Li, H., Consortium, G., 2013. Substitution of Hainan indigenous genetic lineage in the Utsat people, exiles of the Champa kingdom. Journal of Systematics and Evolution 51(3), 287–294. https://doi.org/10.1111/jse.12000.

Lipman, J.N., 1998. Familiar strangers: a history of Muslims in Northwest China. Hong Kong University Press.

Liu, S., Huang, S., Chen, F., Zhao, L., Yuan, Y., Francis, S.S., Fang, L., Li, Z., Lin, L., Liu, R., Zhang, Y., Xu, H., Li, S., Zhou, Y., Davies, R.W., Liu, Q., Walters, R.G., Lin, K., Ju, J., Korneliussen, T., Yang, M.A., Fu, Q., Wang, J., Zhou, L., Krogh, A., Zhang, H., Wang, W., Chen, Z., Cai, Z., Yin, Y., Yang, H., Mao, M., Shendure, J., Wang, J., Albrechtsen, A., Jin, X., Nielsen, R., Xu, X., 2018. Genomic Analyses from Non-invasive Prenatal Testing Reveal Genetic Associations, Patterns of Viral Infections, and Chinese Population History. Cell 175(2), 347–359 e314. https://doi.org/10.1016/j.cell.2018.08.016.

Liu, Y., Yang, J., Li, Y., Tang, R., Yuan, D., Wang, Y., Wang, P., Deng, S., Zeng, S., Li, H., Chen, G., Zou, X., Wang, M., He, G., 2021a. Significant East Asian Affinity of the Sichuan Hui Genomic Structure Suggests the Predominance of the Cultural Diffusion Model in the Genetic Formation Process. Front Genet 12, 626710. https://doi.org/10.3389/fgene.2021.626710.

Liu, Y., Yang, J., Li, Y., Tang, R., Yuan, D., Wang, Y., Wang, P., Deng, S., Zeng, S., Li, H., Chen, G., Zou, X., Wang, M., He, G., 2021b. Significant East Asian Affinity of the Sichuan Hui Genomic Structure Suggests the Predominance of the Cultural Diffusion Model in the Genetic Formation Process. Front Genet 12(834), 626710. https://doi.org/10.3389/fgene.2021.626710.

Loh, P.R., Lipson, M., Patterson, N., Moorjani, P., Pickrell, J.K., Reich, D., Berger, B., 2013. Inferring admixture histories of human populations using linkage disequilibrium. Genetics 193(4), 1233–1254. https://doi.org/10.1534/genetics.112.147330.

Ma, X., Yang, W., Gao, Y., Pan, Y., Lu, Y., Chen, H., Lu, D., Xu, S., 2021. Genetic Origins and Sex-Biased Admixture of the Huis. Mol Biol Evol 38(9), 3804–3819. https://doi.org/10.1093/molbev/msab158.

Mao, X., Zhang, H., Qiao, S., Liu, Y., Chang, F., Xie, P., Zhang, M., Wang, T., Li, M., Cao, P., Yang, R., Liu, F., Dai, Q., Feng, X., Ping, W., Lei, C., Olsen, J.W., Bennett, E.A., Fu, Q., 2021. The deep population history of northern East Asia from the Late Pleistocene to the Holocene. Cell 184(12), 3256–3266 e3213. https://doi.org/10.1016/j.cell.2021.04.040.

Meng, H.T., Han, J.T., Zhang, Y.D., Liu, W.J., Wang, T.J., Yan, J.W., Huang, J.F., Du, W.A., Guo, J.X., Wang, H.D., 2014. Diversity study of 12 Xūchromosomal STR loci in H ui ethnic from C hina. Electrophoresis 35(14), 2001–2007.

Ning, C., Li, T., Wang, K., Zhang, F., Li, T., Wu, X., Gao, S., Zhang, Q., Zhang, H., Hudson, M.J., Dong, G., Wu, S., Fang, Y., Liu, C., Feng, C., Li, W., Han, T., Li, R., Wei, J., Zhu, Y., Zhou, Y., Wang, C.C., Fan, S., Xiong, Z., Sun, Z., Ye, M., Sun, L., Wu, X., Liang, F., Cao, Y., Wei, X., Zhu, H., Zhou, H., Krause, J., Robbeets, M., Jeong, C., Cui, Y., 2020. Ancient genomes from northern China suggest links between subsistence changes and human migration. Nat Commun 11(1), 2700. https://doi.org/10.1038/s41467-020-16557-2.

Olson, J.S., 1998. An ethnohistorical dictionary of China. Greenwood Publishing Group.

Patterson, N., Moorjani, P., Luo, Y., Mallick, S., Rohland, N., Zhan, Y., Genschoreck, T., Webster, T., Reich, D., 2012. Ancient admixture in human history. Genetics 192(3), 1065–1093. https://doi.org/10.1534/genetics.112.145037.

Patterson, N., Price, A.L., Reich, D., 2006. Population structure and eigenanalysis. PLoS Genet 2(12), e190. https://doi.org/10.1371/journal.pgen.0020190.

Pickrell, J.K., Pritchard, J.K., 2012. Inference of population splits and mixtures from genome-wide allele frequency data. PLoS Genet 8(11), e1002967. https://doi.org/10.1371/iournal.pgen.1002967.

Thurgood, G., 2006. Sociolinguistics and Contact-induced Language Change: Hainan Cham, Anong, and Phan Rang Cham. Tenth International Conference on Austronesian Linguistics, 17-20 January 2006, Palawan, Philippines. Linguistic Society of the Philippines and SIL International.

Tinker, N.A., Mather, D.E., 1993. Kin - Software for Computing Kinship Coefficients. Journal of Heredity 84(3), 238–238. https://doi.org/10.1093/oxfordjournals.jhered.a111330.

Tran, N.T., Reid, A., 2006. Viêt Nam: borderless histories. Univ of Wisconsin Press.

Wang, C.C., Lu, Y., Kang, L., Ding, H., Yan, S., Guo, J., Zhang, Q., Wen, S.Q., Wang, L.X., Zhang, M., Tong, X., Huang, X., Nie, S., Deng, Q., Zhu, B., Jin, L., Li, H., 2019. The massive assimilation of indigenous East Asian populations in the origin of Muslim Hui people inferred from paternal Y chromosome. Am J Phys Anthropol 169(2), 341–347. https://doi.org/10.1002/aipa.23823.

Wang, C.C., Yeh, H.Y., Popov, A.N., Zhang, H.Q., Matsumura, H., Sirak, K., Cheronet, O., Kovalev, A., Rohland, N., Kim, A.M., Mallick, S., Bernardos, R., Tumen, D., Zhao, J., Liu, Y.C., Liu, J.Y., Mah, M., Wang, K., Zhang, Z., Adamski, N., Broomandkhoshbacht, N., Callan, K., Candilio, F., Carlson, K.S.D., Culleton, B.J., Eccles, L., Freilich, S., Keating, D., Lawson, A.M., Mandl, K., Michel, M., Oppenheimer, J., Ozdogan, K.T., Stewardson, K., Wen, S., Yan, S., Zalzala, F., Chuang, R., Huang, C.J., Looh, H., Shiung, C.C., Nikitin, Y.G., Tabarev, A.V., Tishkin, A.A., Lin, S., Sun, Z.Y., Wu, X.M., Yang, T.L., Hu, X., Chen, L., Du, H., Bayarsaikhan, J., Mijiddorj, E., Erdenebaatar, D., Iderkhangai, T.O., Myagmar, E., Kanzawa-Kiriyama, H., Nishino, M., Shinoda, K.I., Shubina, O.A., Guo, J., Cai, W., Deng, Q., Kang, L., Li, D., Li, D., Lin, R., Nini, Shrestha, R., Wang, L.X., Wei, L., Xie, G., Yao, H., Zhang, M., He, G., Yang, X., Hu, R., Robbeets, M., Schiffels, S., Kennett, D.J., Jin, L., Li, H., Krause, J., Pinhasi, R., Reich, D., 2021. Genomic insights into the formation of human populations in East Asia. Nature 591(7850), 413–419. https://doi.org/10.1038/s41586-021-03336-2.

Wang, Q., Zhao, J., Ren, Z., Sun, J., He, G., Guo, J., Zhang, H., Ji, J., Liu, Y., Yang, M., Yang, X., Chen, J., Zhu, K., Wang, R., Li, Y., Chen, G., Huang, J., Wang, C.C., 2020a. Male-Dominated Migration and Massive Assimilation of Indigenous East Asians in the Formation of Muslim Hui People in Southwest China. Front Genet 11(1742), 618614. https://doi.org/10.3389/fgene.2020.618614.

Wang, Q., Zhao, J., Ren, Z., Sun, J., He, G., Guo, J., Zhang, H., Ji, J., Liu, Y., Yang, M., Yang, X., Chen, J., Zhu, K., Wang, R., Li, Y., Chen, G., Huang, J., Wang, C.C., 2020b. Male-Dominated Migration and Massive Assimilation of Indigenous East Asians in the Formation of Muslim Hui People in Southwest China. Front Genet 11, 618614. https://doi.org/10.3389/fgene.2020.618614.

Wang, T., Wang, W., Xie, G., Li, Z., Fan, X., Yang, Q., Wu, X., Cao, P., Liu, Y., Yang, R., Liu, F., Dai, Q., Feng, X., Wu, X., Qin, L., Li, F., Ping, W., Zhang, L., Zhang, M., Liu, Y., Chen, X., Zhang, D., Zhou, Z., Wu, Y., Shafiey, H., Gao, X., Curnoe, D., Mao, X., Bennett, E.A., Ji, X., Yang, M.A., Fu, Q., 2021. Human population history at the crossroads of East and Southeast Asia since 11,000 years ago. Cell 184(14), 3829–3841 e3821. https://doi.org/10.1016/j.cell.2021.05.018.

Wu, D., Dou, J., Chai, X., Bellis, C., Wilm, A., Shih, C.C., Soon, W.W.J., Bertin, N., Lin, C.B., Khor, C.C., DeGiorgio, M., Cheng, S., Bao, L., Karnani, N., Hwang, W.Y.K., Davila, S., Tan, P., Shabbir, A., Moh, A., Tan, E.K., Foo, J.N., Goh, L.L., Leong, K.P., Foo, R.S.Y., Lam, C.S.P., Richards, A.M., Cheng, C.Y., Aung, T., Wong, T.Y., Ng, H.H., Consortium, S.K., Liu, J., Wang, C., 2019. Large-Scale Whole-Genome Sequencing of Three Diverse Asian Populations in Singapore. Cell 179(3), 736–749 e715. https://doi.org/10.1016/j.cell.2019.09.019.

Xie, M., Song, F., Li, J., Lang, M., Luo, H., Wang, Z., Wu, J., Li, C., Tian, C., Wang, W., Ma, H., Song, Z., Fan, Y., Hou, Y., 2019. Genetic substructure and forensic characteristics of Chinese Hui populations using 157 Y-SNPs and 27 Y-STRs. Forensic Sci Int Genet 41, 11–18. https://doi.org/10.1016/j.fsigen.2019.03.022.

Xie, T., Guo, Y., Chen, L., Fang, Y., Tai, Y., Zhou, Y., Qiu, P., Zhu, B., 2018. A set of autosomal multiple InDel markers for forensic application and population genetic analysis in the Chinese Xinjiang Hui group. Forensic Science International: Genetics 35, 1–8.

Xu, S., Yin, X., Li, S., Jin, W., Lou, H., Yang, L., Gong, X., Wang, H., Shen, Y., Pan, X., He, Y., Yang, Y., Wang, Y., Fu, W., An, Y., Wang, J., Tan, J., Qian, J., Chen, X., Zhang, X., Sun, Y., Zhang, X., Wu, B., Jin, L., 2009. Genomic dissection of population substructure of Han Chinese and its implication in association studies. Am J Hum Genet 85(6), 762–774. https://doi.org/10.1016/j.ajhg.2009.10.015.

Yang, M.A., Fan, X., Sun, B., Chen, C., Lang, J., Ko, Y.C., Tsang, C.H., Chiu, H., Wang, T., Bao, Q., Wu, X., Hajdinjak, M., Ko, A.M., Ding, M., Cao, P., Yang, R., Liu, F., Nickel, B., Dai, Q., Feng, X., Zhang, L., Sun, C., Ning, C., Zeng, W., Zhao, Y., Zhang, M., Gao, X., Cui, Y., Reich, D., Stoneking, M., Fu, Q., 2020. Ancient DNA indicates human population shifts and admixture in northern and southern China. Science 369(6501), 282–288. https://doi.org/10.1126/science.aba0909.

Yang, X., Zhang, X., Zhu, J., Chen, L., Liu, C., Feng, X., Chen, L., Wang, H., Liu, C., 2017. Genetic analysis of 19 X chromosome STR loci for forensic purposes in four Chinese ethnic groups. Sci. Rep. 7(1), 1–11.

Yao, H., Wang, M., Zou, X., Li, Y., Yang, X., Li, A., Yeh, H.Y., Wang, P., Wang, Z., Bai, J., Guo, J., Chen, J., Ding, X., Zhang, Y., Lin, B., Wang, C.C., He, G., 2021. New insights into the fine-scale history of western-eastern admixture of the northwestern Chinese population in the Hexi Corridor via genome-wide genetic legacy. Molecular genetics and genomics: MGG 296(3), 631–651. https://doi.org/10.1007/s00438-021-01767-0.

Yao, H.B., Wang, C.C., Tao, X., Shang, L., Wen, S.Q., Zhu, B., Kang, L., Jin, L., Li, H., 2016. Genetic evidence for an East Asian origin of Chinese Muslim populations Dongxiang and Hui. Sci Rep 6, 38656. https://doi.org/10.1038/srep38656.

Yao, Y.-G., Kong, Q.-P., Wang, C.-Y., Zhu, C.-L., Zhang, Y.-P., 2004. Different matrilineal contributions to genetic structure of ethnic groups in the silk road region in china. Molecular biology and evolution 21(12), 2265–2280.

Zhao, J., Wurigemule, Sun, J., Xia, Z., He, G., Yang, X., Guo, J., Cheng, H.Z., Li, Y., Lin, S., Yang, T.L., Hu, X., Du, H., Cheng, P., Hu, R., Chen, G., Yuan, H., Zhang, X.F., Wei, L.H., Zhang, H.Q., Wang, C.C., 2020. Genetic substructure and admixture of Mongolians and Kazakhs inferred from genome-wide array genotyping. Ann Hum Biol 47(7-8), 620–628. https://doi.org/10.1080/03014460.2020.1837952.

Zhao, Q., Bian, Y., Zhang, S., Zhu, R., Zhou, W., Gao, Y., Li, C., 2017. Population genetics study using 26 Y-chromosomal STR loci in the Hui ethnic group in China. Forensic Sci Int Genet 28, e26–e27. https://doi.org/10.1016/j.fsigen.2017.01.018.

Zhou, B., Wen, S., Sun, H., Zhang, H., Shi, R., 2020. Genetic affinity between Ningxia Hui and eastern Asian populations revealed by a set of InDel loci. Royal Society open science 7(1), 190358.

Zhou, Y., Zhou, B., Pache, L., Chang, M., Khodabakhshi, A.H., Tanaseichuk, O., Benner, C., Chanda, S.K., 2019. Metascape provides a biologist-oriented resource for the analysis of systems-level datasets. Nat Commun 10(1), 1523. https://doi.org/10.1038/s41467-019-09234-6.

Zou, X., Wang, Z., He, G., Wang, M., Liu, J., Wang, S., Ye, Z., Wang, F., Hou, Y., 2020. Genetic variation and population structure analysis of Chinese Wuzhong Hui population using 30 Indels. Ann. Hum. Biol. 47(3), 300–303. https://doi.org/10.1080/03014460.2020.1736627.

